# High-frequency amplitude-modulated sinusoidal stimulation desynchronizes neural activity and enhances naturalness of evoked sensations

**DOI:** 10.1101/2024.02.14.580219

**Authors:** Beatrice Barra, David Samuel Rose, Ritesh Kumar, Chaitanya Gopinath, Ehsan Mirzakhalili, Scott F. Lempka, Robert A. Gaunt, Eric Daniel Głowacki, Lee E Fisher

**Author notes:** Corresponding author: Lee E. Fisher.

## Abstract

Regaining sensory feedback is critical for people living with limb amputation. Electrical stimulation of sensory fibers in peripheral nerves has been shown to restore focal percepts in the missing limb. However, conventional rectangular current pulses induce sensations often described as unnatural, likely due to the synchronous and periodic nature of activity they evoke. Here we introduce a fast-oscillating amplitude-modulated sinusoidal (FAMS) stimulation waveform that desynchronizes neural activity. Using computational modeling, we show that sinusoidal waveforms evoke asynchronous and irregular firing patterns, with frequency-dependent effects. Leveraging both low- and high-frequency mechanisms, FAMS exploits membrane nonlinearities to enhance neuron-specific differences. In a feline model of peripheral nerve stimulation, FAMS evoked activity that was more asynchronous than conventional rectangular pulses, while remaining easily controllable with simple stimulation parameters. Importantly, in human experiments using noninvasive stimulation of the median nerve, participants reported that FAMS evoked more natural sensations compared to rectangular biphasic pulses in a two-alternative forced-choice task. The preference for FAMS increased at higher intensities above sensory threshold, despite intensity-matched stimulation across waveform types. These findings provide evidence that reduced synchrony in afferent recruitment translates into more naturalistic and comfortable sensory percepts. Together, our results establish FAMS as a promising biomimetic stimulation strategy with potential for clinical applications in sensory feedback restoration.

**One Sentence Summary:** A new electrical stimulation waveform allows for evoking and controlling more naturalistic neural activity than can be achieved with traditional stimulation waveforms.

## INTRODUCTION

Electrical stimulation has been used to restore physiological autonomic, sensory, and motor functions following a wide spectrum of neurological and traumatic disorders, such as paralysis^1–7^, stroke^8,9^, limb amputation^10–19^, and chronic pain^20–23^. Traditionally, electrical stimulation paradigms have relied on charge-balanced rectangular pulses. Rectangular pulses have been shown to be safe and effective ^24^ and currently represent the basic unit of stimulation. In most applications, a train of pulses is delivered with a specific constant stimulation frequency. The number of activated fibers is determined by the charge injected by each pulse, which is intrinsically determined by the specific combination of the amplitude and duration of the pulse^24^. With this approach, all recruited fibers fire synchronously, with each stimulus pulse evoking a single simultaneous action potential volley in all responding neurons^25,26^.

Prior studies have hypothesized that highly synchronous firing patterns, especially those with highly consistent inter-pulse timing, are responsible for several negative effects of electrical stimulation, such as the production of unnatural paresthetic sensory percepts^10,12,27,28^ in people with limb amputation, and rapid muscle fatigue during functional electrical stimulation in people with spinal cord injury or stroke^29–33^. To overcome these limitations, recent studies have pursued the design of *biomimetic* stimulation protocols that aim to improve the naturalness of stimulation-induced neural activity by emulating natural neural activity. These biomimetic approaches typically take one of two forms. In one form, the pulse train is designed to mimic the time-varying firing rate of specific neurons (e.g., rapidly adapting cutaneous afferents) by manually tuning the inter-stimulus intervals throughout the stimulation train^10,17,34–36^. This type of approach aims at inducing more naturalistic activity within each fiber than traditional constant frequency trains^37^, by incorporating a limited level of stochasticity in the firing patterns of individual neurons^38,39^ . However, activity is still induced synchronously in all recruited neurons and as such, downstream neurons and muscles still receive highly synchronized input, which may result in non-natural activity in those targets. A second form of biomimetic stimulation utilizes amplitude-modulated trains of rectangular pulses at kHz frequencies^26,40^. This approach achieves partial desynchronization of neural activity across subpopulations, and could therefore be very useful for motor applications: some degree of synchronization within a specific population would not only be acceptable, but also useful to provide strong muscle contractions^40^. Multiple recent studies have sought to characterize the effect of high-frequency rectangular pulses on the quality of evoked sensory percepts. While some have shown that the use of transcutaneous high-frequency stimulation of the spinal cord does not impact the quality of evoked sensory percepts^41^, others have suggested that during continuous and unmodulated high-frequency stimulation, the high degree of desynchronization across neurons might prevent the generation of paresthesia due to transmission failure to the somatosensory cortex^28^.

Overall, all of the above-mentioned approaches rely on the use of rectangular pulses, which inevitably generate synchronous firing across all neurons that are recruited by each pulse. In fact, during a rectangular stimulation pulse, neurons near the stimulation electrode exhibit impulse-responses that depend on their membrane physiological properties as well as their geometric structure^42^. In a homogeneous population of neurons, the similarity of membrane properties causes neurons near the electrode to respond synchronously to the pulse^28^. This is especially true for stimulation of peripheral nerves, in which axons for many neurons run parallel to each other along the nerve. In these circumstances, desynchronization is solely achieved by delivering stimulation pulses quickly enough to leverage the refractory period of responding neurons^28^. As a consequence, methods based on square pulses are unable to modulate firing rates in a controlled fashion while also achieving substantial population-level desynchronization.

Here, we address this problem by designing a sinusoidal stimulation waveform that induces asynchronous, and quasi-stochastic neural activity, while also allowing specific stimulation parameters to modulate and control firing rate profiles. We hypothesized that by using sinusoidal waveforms to induce a more gradual charge injection we can enhance the desynchronization of neural activity in a neural population. We used a combined computational modeling and experimental approach to investigate the strengths and weaknesses of low-frequency and high-frequency sinusoidal stimulation. We then designed a novel fast amplitude modulated sinusoidal waveform (FAMS), consisting of a kHz carrier frequency sinusoid that is amplitude-modulated at a lower frequency. We hypothesized that this waveform would combine the positive effects of low- and high-frequency sinusoids, while avoiding unwanted effects of unmodulated low- or high-frequency sinusoids (i.e., higher injected charge, conduction block)^43,44^. In our combined computational modeling and experimental approach, we show that FAMS induces quasi-stochastic firing at the single neuron level and enhances desynchronization across fibers. We investigated the mechanisms that induce asynchronous and aperiodic firing and showed that FAMS-evoked neural activity can be modulated and controlled robustly with a small set of parameters. We then used FAMS to transcutaneously stimulate the median nerve of human subjects and found that FAMS is perceived as more natural than rectangular-pulse electrical stimulation.

## RESULTS

### Sinusoidal stimulation desynchronizes firing in a simulated neural population

We hypothesized that sinusoidal stimulation would generate a smoother charge injection profile than rectangular pulses, thus enhancing desynchronization of neural activity within a population. To test this hypothesis, we used a biophysical computational model (**Figure 1A**) to compare the response of neurons to rectangular pulses and a sinusoidal stimulation waveform. We represented our stimulating electrode as a pair of ideal point sources. We then calculated the corresponding electric potential fields using the superposition principle under a quasistatic approximation^45^. We then simulated a population of axons (number ranging between 55 and 65) in a space of 100 μm^2^ using the NEURON simulation environment^46^ (see Methods). We simulated a heterogenous population of sensory neurons by varying neurons’ diameters between 6 and 12 μm. **Figure 1B** shows the response of this population to a series of rectangular pulses and a continuous sinusoid, each delivered at a current amplitude of 1.5 times its respective excitation threshold. In both cases, neurons tend to exhibit periodic firing behavior, where they fire shortly after the onset of the cathodic phase of stimulation. We quantified this effect by computing a peri-stimulus time histogram (PSTH; i.e., calculating the time it took each neuron to fire an action potential after the onset of stimulation). For the rectangular pulse, post-stimulus spike time was computed with respect to the beginning of the cathodic phase of the pulse (**Figure 1B**, (ii), top panel); for the sinusoid, post-stimulus spike time was computed with respect to the beginning of the cathodic phase of the sinusoid cycle (**Figure 1B** (ii) bottom panel). **Figure 1B** (iii) shows PSTHs in both conditions. Indeed, sinusoidal stimulation evoked variable firing patterns with spike timings that span the whole duration of the cathodic phase and were far more variable than the patterns evoked by rectangular pulses. Interestingly, neurons fired both during the cathodic and anodic phases of the sinusoidal waveform, likely because we used a bipolar stimulation configuration. These results corroborate the hypothesis that a sinusoidal stimulation profile could enhance firing desynchronization across a neuronal population. We next explored how the frequency of sinusoidal waveforms influences this effect to determine which stimulation frequencies would most efficiently achieve desynchronization.

**Figure 1.**
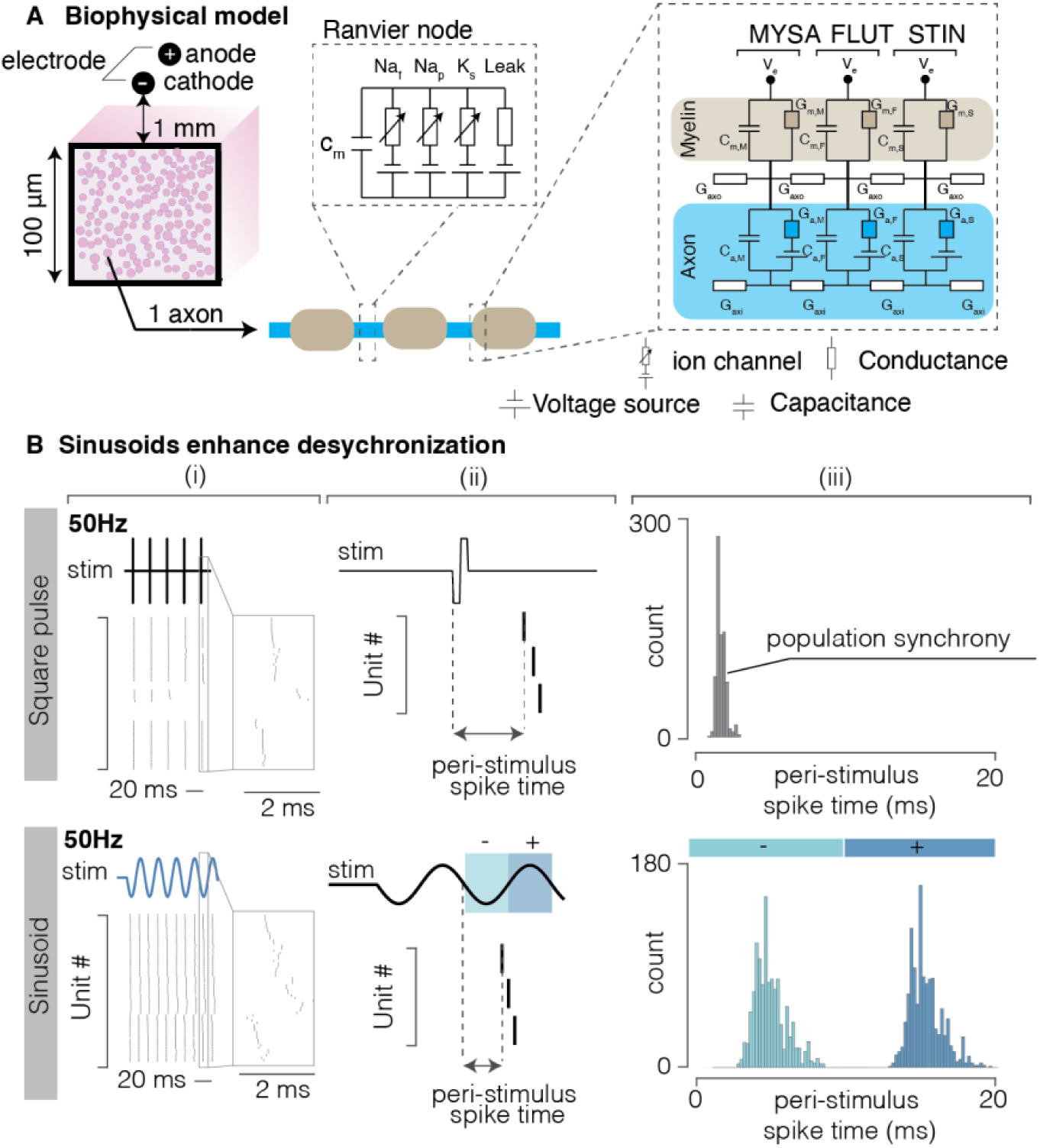
Computational Model. **(A)** Schematic of simulated neuronal population. Myelinated axons (not to scale) are placed in a limited volume (100 um^2^) and adjacent to a pair of electrodes (- for cathode and + for anode). The electrodes are 1 mm apart and are placed 1 mm from the neurons, parallel to the axial direction of the axons. On the right, the myelinated region is explicitly divided in the segments MYSA i.e. the myelin attachment segment, FLUT i.e. the paranode main segment and STIM, i.e. the internode segment. The nodes of Ranvier include slow potassium channels, persistent and fast potassium channels, as well as leakage current. **(B)**, Computational simulation of the effect of single pulse stimulation compared to sinusoidal waveforms. Top row, rectangular pulses (50 Hz). Bottom row, sinusoidal stimulation (50 Hz). (i) Raster of spikes evoked by both types of stimulation. The sinusoidal stimulation displays a noticeable time-dispersion of spiking activity, compared to the rectangular pulse. (ii) Schematic illustrating how the post-stim spike time has been computed for the rectangular pulses and sinusoid. (iii) Peri-stimulus time histogram of spiking evoked by the two stimulation waveforms.

### Desynchronization via sinusoidal stimulation is frequency-dependent

Neurons typically fire one action potential in response to a single rectangular stimulus pulse. Therefore, provided that the pulse frequency is low enough to allow the neuron to fully recover from its refractory period, the frequency of stimulation directly determines the firing rate of the activated neurons. Conversely, variations in the frequency of sinusoidal waveforms intrinsically changes the timescale of charge injection in the extracellular medium. We therefore hypothesized that the frequency of sinusoidal stimulation would impact the firing behavior of neurons in a complex manner, both at the population and the single neuron level. First, we used our computational model to characterize the effects of stimulation frequency on firing desynchronization of the neuronal population. We delivered sinusoidal stimulation at 50, 100, 500, and 1,000 Hz and analyzed the PSTH in each condition. For each frequency, we used a current amplitude value equal to 1.5 times the stimulation threshold. Lower frequencies exhibited a higher degree of desynchronization (**Figure 2A, 2C**) which gradually decreased with increasing frequency. The marked bimodal nature of the PSTH at lower frequencies (corresponding to responses to both the anodic and cathodic phases of the waveform) was lost at higher frequencies, likely because in this regime, the anodic phase of the stimulation occurs during the refractory period of action potentials generated during the preceding cathodic phase, hence hindering the generation of another action potential.

**Figure 2.**
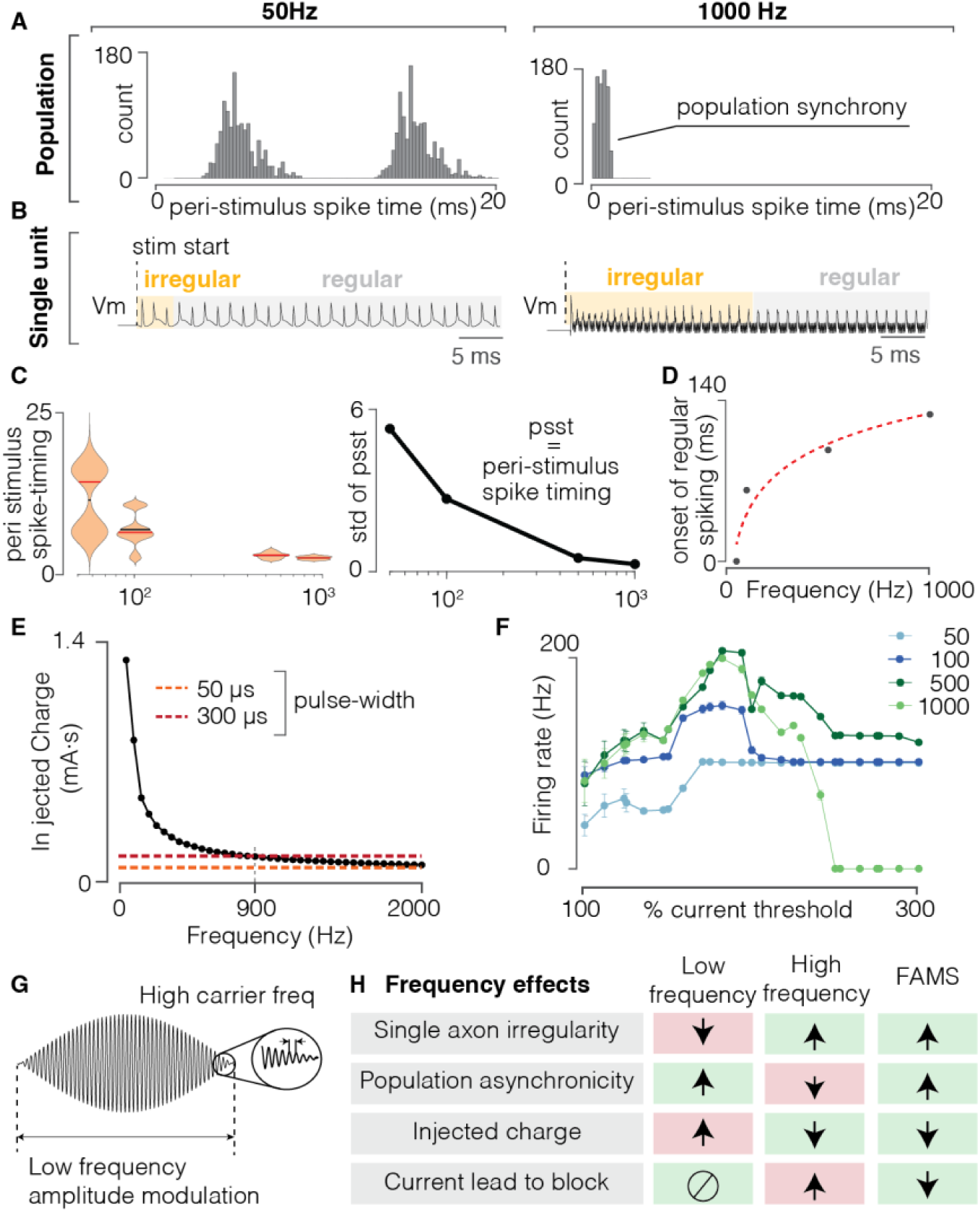
Sinusoidal stimulation elicits asynchronous and quasi-stochastic activity that is frequency dependent. **(A)** Peri-stimulus time histogram of spiking evoked by the sinusoidal stimulation at 50 Hz (left) and 1000 Hz (right) in a neural population. **(B)** Representation of evoked spiking activity in a simulated single axon: yellow shadowed area indicates the duration of irregular spiking, the grey shaded area indicates the time at which the neuron displayed regular spiking activity. **(C)** violin plot of post-stim spike time distributions for sinusoidal stimulation at different frequencies. Red: median value of the distribution. Black line: mean value of the distribution. On the right, quantification of the standard deviation of the peri-stimulus spike time for each distribution. **(D)** time of onset of regular spiking patterns, as a function of sinusoid frequency. Higher frequencies generate irregular spiking for more extended periods of time. Dashed red line: exponential fit of datapoints, y = a·x^b^ + c (a = -2954, b = -0.8, c = 130.1, R^2^ = 0.968). **(E)** For sinusoidal stimulation, value of injected charge during cathodic phase of a sinusoid as a function of frequency. Orange dotted line: injected charge for a rectangular pulse of pulse-width equal to 50 μs; Red dashed line: injected charge for a rectangular pulse of pulse-width equal to 300 μs. Current values are fixed at 1.2 times the threshold amplitude, for each condition. **(F)** Average firing rate as a function of increasing stimulation amplitude, for sinusoids at different frequencies (color code: light blue, 50Hz; dark blue, 100Hz; dark green, 500 Hz, light green, 1000Hz). Current amplitude values are expressed as percentages of 1.2 times the threshold value, where threshold was defined as the current needed to elicit at least two spikes. **(G)** Illustration of the FAMS waveform, that consists of a high (i.e., carrier) frequency sinusoid, whose amplitude is modulated at a lower (i.e., beat) frequency. **(H)** Schematic illustrating advantages and disadvantages of low and high- frequency sinusoidal stimulation, as well as FAMS stimulation. We hypothesized that by combining a low frequency and high frequency component, FAMS would optimize neural recruitment to generate asynchronous and stochastic firing in a neural population

We next analyzed the effect of waveform frequency at the single axon level (**Figure 2B**). At each of the explored frequencies (i.e. 50, 100, 500, and 1,000 Hz), the transmembrane voltage displayed firing behavior that was initially irregular, exhibiting variations in inter-spike intervals and spike peak amplitude. This period of irregular firing was then followed by a very regular firing behavior, with repeated inter-spike intervals and constant spike amplitude. The duration of this irregular firing behavior increased with increasing waveform frequency (**Figure 2B, 2D**). We calculated the duration of the irregular regime and computed the phase-difference (**Figure S1**) between the stimulus and the generation of action potentials during the irregular and regular firing period. Regular firing was associated with a stable phase lag between stimulus and transmembrane voltage at the instants preceding the action potentials (**Figure S1A**). Conversely, irregular firing was associated with a higher variability in the phase differences between the stimulus and the membrane potential. We argue that this transitory irregular firing behavior in response to high-frequency waveforms could be leveraged to enhance irregularity and desynchronization in population firing patterns.

In summary, our results highlight that while a higher frequency enhances irregular firing at the single neuron level, it concurrently increases synchronization of activity across neurons. In addition, previous studies have highlighted that (1) low-frequency sinusoidal waveforms require injection of large amounts of charge (**Figure 2E**), which can cause damage to the electrodes and surrounding biological tissue and (2) high-frequency sinusoids often cause conduction block^43,44^, eventually leading to a halt in firing (**Figure 2F**). These conflicting effects of low and high-frequency waveforms suggest that while simple sinusoidal waveforms represent an improvement over rectangular pulses with respect to asynchronous firing behavior, they are unlikely to achieve naturalistic firing behavior. We therefore hypothesized that a waveform combining low- and high-frequency features would achieve both asynchronous activity at the population level and quasi-stochastic firing at the single neuron level, while avoiding conduction block effects and excessive charge injection. To leverage the advantages of both low- and high-frequency sinusoidal stimulation, we implemented a novel stimulation waveform consisting of a high-frequency sinusoid (1 to 8 kHz) whose amplitude is modulated at a lower frequency (10 to 50 Hz) (**Figure 2G**). Based on the results of our computational model, we hypothesized that the high frequency component would avoid excessive charge injection while fostering irregular firing at the single neuron level. Additionally, the low-frequency modulation would prevent the neurons from entering a conduction block state, while also enhancing desynchronization of firing at the population level (**Figure 2H**). We called this waveform a Fast Amplitude-Modulated Sinusoid (FAMS), described mathematically by equation (1):

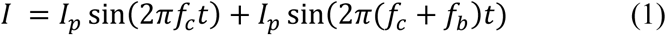

where I_p_ is the peak current amplitude, f_c_ is the high (*carrier*) frequency, and f_b_ is the low (*beat*) frequency.

### Rectification by ion channels causes asynchronous firing during FAMS

We used our computational model to investigate whether the FAMS waveform would generate asynchronous firing in a population of parallel axons. Using the same geometry previously described, we delivered pulses of FAMS at increasing amplitude (1000 Hz carrier frequency and 10 Hz beat frequency) and inspected the firing patterns at the population level. Spiking activity started shortly after the waveform onset and persisted for approximately 80% of the stimulus duration. Most axons displayed asynchronous firing of action potentials (**Figure 3A** (i)). We quantified the overall level of desynchronization by computing the time between each spike and the closest spike for every other neuron in the population. We then averaged this measure to obtain one value associated with each spike (**Figure 3A** (ii)). This desynchronization metric increases both when axons display irregular firing and when population asynchronicity increases. During FAMS, action potentials were on average 3 ms apart (3.0 ± 2.4 ms), which was significantly higher than all other stimulation waveforms (**Figure 3A** (ii)). Interestingly, desynchronization for high-frequency waveforms was higher than for low-frequency, suggesting that the initial axon irregular firing has a lasting impact in generating asynchronous firing at the population level. To explore the mechanisms underlying asynchronous firing during FAMS, we then delivered FAMS to a single 10 µm diameter axon placed at 1 mm from the stimulating electrode. We used a single pulse of sub-threshold FAMS stimulation at increasing amplitudes and observed that higher peak amplitudes generate more asymmetrical transmembrane currents (**Figure 3B**). A spectral analysis of the transmembrane current profiles revealed that at higher stimulation currents, transmembrane current oscillations contained greater low frequency components. This behavior is coherent with a model of demodulation, where a low frequency component arises from input rectification followed by low pass filtering ^44^. As with previous literature, we hypothesized that the stimulus rectification stemmed from non-linearities in the dynamics of membrane ion channels^44^. To test this hypothesis, we independently and systematically modulated the gating time constants of each voltage-gated ion channel by ±50%, by changing their forward and backward rates without affecting their steady-state activation and inspected differences in rectification properties^44^. Changes in the dynamics of any of the gating variables increased or decreased the amount of resulting rectification (**Figure 3C**). This result suggests that the demodulation of the FAMS waveform is due to the gating properties of ion channels. As those gating properties are voltage dependent, we reasoned that the asynchronous firing could arise from slight variations in stimulus amplitude. We verified that the time of first spike during the stimulation waveform decreased with increasing stimulation amplitude (**Figure 3D**, (i)). We reasoned that neurons in a population are bound to receive slightly different stimulation amplitudes due to small differences in locations. Indeed, by systematically changing the position of the axon, we found that time of spiking onset increased with increasing distance from the electrode (**Figure 3D**, (ii)). Similarly, we systematically changed the diameter of the axon and verified that the time of firing also depended on the fiber diameter (**Figure 3D**, (iii)). We concluded that the asynchronous firing in a population of neurons arises as a result of a combination of a neuron’s position and diameter. By inspecting the values of gating variables at spiking initiation we observed that each neuron generated an action potential in correspondence to a distinct channel gating configuration (**Figure 3E**). Those variations also propagated to and even intensified in the following spike, suggesting that asynchronous and irregular firing may be maintained during the whole stimulation waveform.

**Figure 3.**
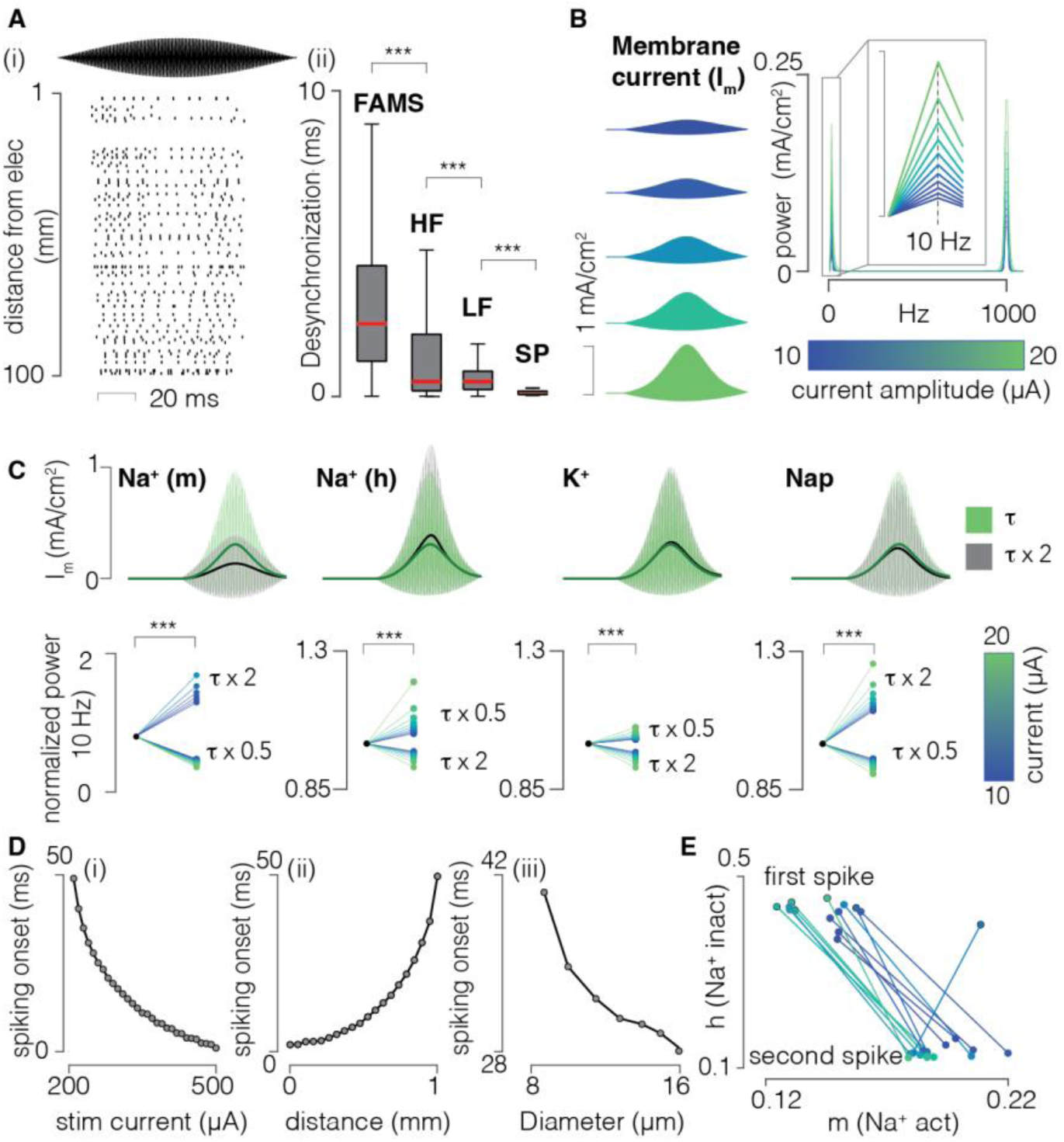
Neuron-specific rectification properties generate asynchronous and quasi-stochastic firing in response to FAMS stimulation. **(A)** Left, raster plot of simulated neurons spiking in response to FAMS stimulation. Neurons fire asynchronously from each other. Right, quantification of the amount of desynchronization for FAMS (carrier frequency 1000Hz, beat frequency 10Hz), high-frequency sinusoids (HF, 1000Hz), low-frequency sinusoids (LF, 50 Hz), and single pulse (SP, 50 Hz). Red line: median value. Outliers not shown. *** p = 0.001, Wilcoxon rank sum test. **(B)** Membrane current profiles at subthreshold stimulus amplitudes, increasing from top to bottom. On the right, the power spectrum of the membrane currents shows both a low- and a high-frequency component. The power spectrum was computed with the fast Fourier transform (FFT). The inset shows that the amount of low-frequency content (peak value at 10 Hz) depends on the stimulus amplitude. **(C)** We increased and decreased the gating variable time constants by ±50**%** by changing the forward and backward rates of each ion channel type. Modified dynamics of each gating variable had an effect on the amount of stimulus rectification which was also dependent of the stimulus amplitude. Top row, membrane current profiles with regular or increased gating variable time constants, for each channel. Bottom row, normalized power at 10 Hz for membrane currents with increased or decreased gating variable time constants. Color represents stimulus current values, blue to green for low to high. *** p = 0.001, Wilcoxon rank sum test. **(D)** Time of spiking onset during the FAMS stimulus, as a function of current amplitude (i), distance from the electrode (ii), and axon’s diameter (iii). Spiking time decreases with increasing current, increases with electrode distance, and decreases with axon diameter. (**E)** Different FAMS amplitudes evoke spiking at different membrane configurations. Dots represent values of Na^+^ gating variables at the time of spiking: Na^+^ activation (m), on the x-axis and Na^+^ inactivation (h), on the y-axis. The specific configuration of sodium channels at the time of spiking depends on the amplitude of FAMS and influences the sodium channels configuration of the following spikes, therefore propagating the asynchronous firing behavior to also affect the second spike.

### FAMS achieves quasi-stochastic firing and population desynchronization in vivo

We then tested FAMS in-vivo to characterize the resulting neural activity. In three cats, we implanted a multi-contact cuff on the sciatic nerve to deliver electrical stimulation while we recorded neural activity from two 32-channel multi-electrode arrays (Blackrock Microsystems,

Salt Lake City, UT, USA) implanted in the L6 and L7 dorsal root ganglia (DRG) (**Figure 4A**). Our goal was to compare the evoked spiking activity in sensory neurons of a peripheral nerve during FAMS and during rectangular pulses, both at the population level as well as in terms of the spike timing in each neuron. We were able to detect single spikes during FAMS but not during rectangular spike stimulation, which generated a compound action potential (CAP). Therefore, to quantify the degree of desynchronization across neurons in the recruited population, we used different methods that depended on the stimulation technique. During FAMS, we quantified desynchronization by computing the time between a selected spike and the closest spike for every other neuron (**Figure 4B**). We then averaged this measure to obtain one value associated with each spike and we repeated this procedure for each occurring spike. For rectangular-pulse stimulation, we computed a similar metric for the CAP response recorded on each DRG microelectrode, comparing the beginning of the evoked response for each electrode to the end of the evoked response for every other electrode (i.e., the most conservative measure of desynchronization) (**Figure S2A**). We compared desynchronization during FAMS (1000 Hz carrier, 10 Hz beat and 100 µA current) and rectangular pulses (10 Hz, 100 µA current) delivered at the same current amplitude. FAMS exhibited substantially higher desynchronization as compared to rectangular pulses in all three animals (+154%, +124%, +91%, for Ct-Fi, Ct-Lu, Ct-Re, respectively, **Figure 4C**). Indeed, the whole distribution of desynchronization measures was considerably higher than the maximum desynchronization achieved with rectangular pulse stimulation (gray box). We computed the dependency of desynchronization on carrier frequency and current amplitude and we found that specific combinations of parameters yield better desynchronization results (**Figure 4D** (i)). We found that those parameter combinations were associated with lower firing rates (**Figure 4D**, (ii)). This result can be intuitively understood by considering that higher firing rates in each neuron will necessarily increase the probability of each neuron spiking at a specific time point. This inversely proportional relationship between desynchronization and firing rate is also present in natural neuronal activity induced by leg movements (**Figure S2C**) and therefore represents a desirable physiological effect.

**Figure 4.**
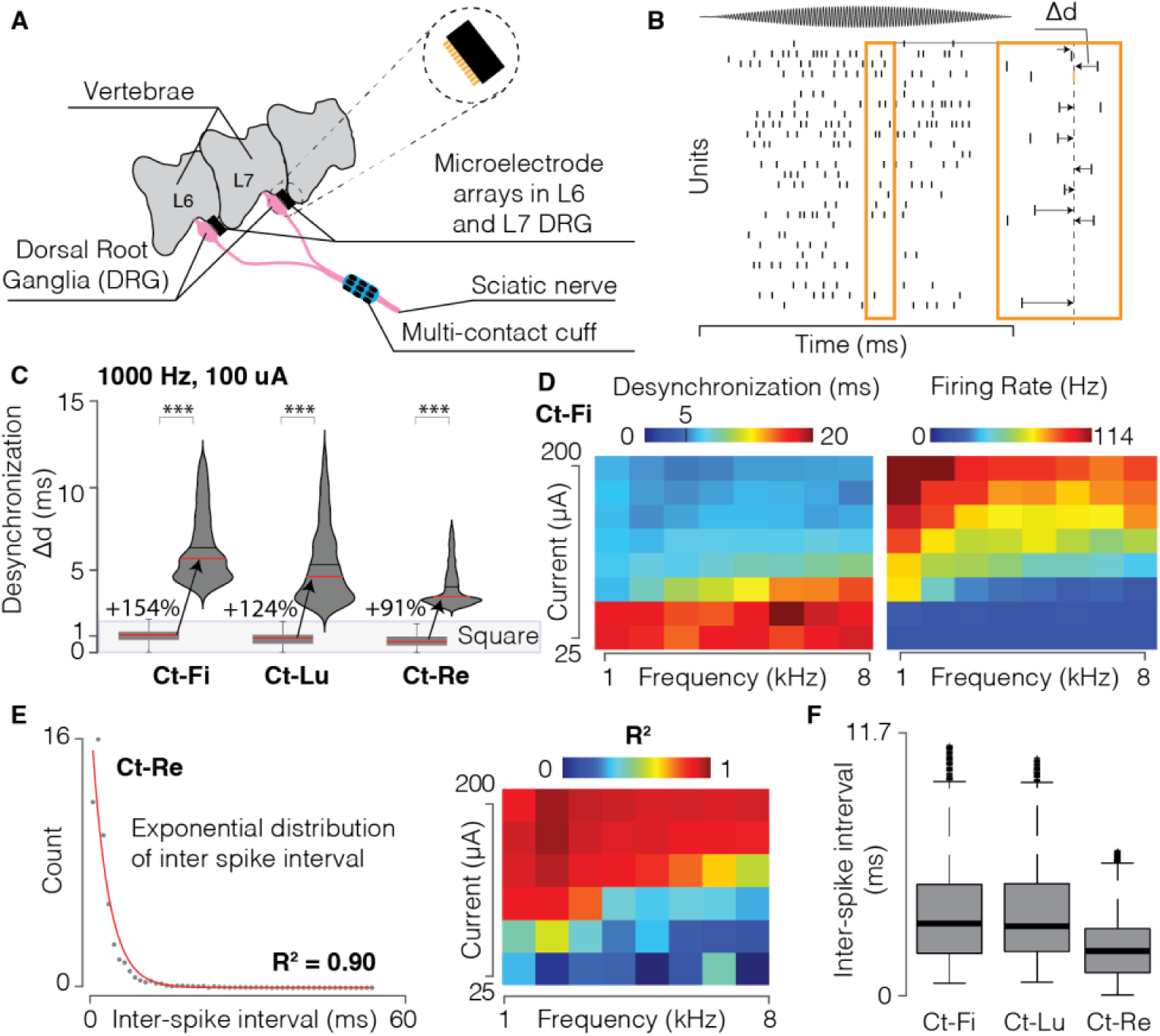
FAMS stimulation elicits asynchronous and quasi-stochastic firing in vivo. **(A)** Schematic of the experimental setup. We implanted a multi-contact cuff-electrode around the sciatic nerve, which we used to deliver electrical stimulation. We used two microelectrode arrays to record neural signals in the dorsal root ganglia (DRG) at the L6 and L7 spinal levels in a feline model. **(B)** Raster plot of spiking activity in neuronal population recorded from the L6 electrode in Ct-Re (1,000 Hz carrier frequency, 75 μA peak current, 10 Hz beat frequency). On the right, schematic of how we calculated desynchronization. For a specific spike (in orange), we computed the time distance from all closest spikes from other neurons and averaged across all computed values. We repeated this measure, for each spike in each neuron. **(C)** Violin plot showing the distribution of desynchronization, computed as explained in (B), for the three animals (Ct-Fi, Ct-Lu, Ct-Re). Distributions are shown in comparison to the desynchronization values obtained during rectangular pulse stimulation (boxplot below). Red line: median value. Black line: mean value. *** p = 0.001, Wilcoxon rank sum test. **(D)** Heatmap of mean desynchronization and mean firing rate values for all current amplitudes and carrier frequencies. **(E)** Left: scatter plot of the inter-spike interval histogram for Ct-Re (1,000 Hz carrier frequency, 75 μA peak current, 10 Hz beat frequency), along with an exponential fit (red line, R^2^ = 0.89). Right: heatmap of R^2^ values for exponential fit of ISI histogram at all parameters combination. An exponential distribution of ISI is indicative of stochastic firing. The R^2^ matrix indicates that quasi-stochastic firing is obtained for all parameter combinations that elicit sustained firing. **(F)** Boxplot of all inter-spike interval values for n=3 animals (Ct-Fi, Ct-Lu, Ct-Re).

We next investigated within-neuron firing patterns during FAMS. Under normal physiological conditions, the firing of each neuron is highly irregular and approximates a stochastic process^38,39,47,48^, exhibiting an exponential distribution of inter-spike intervals (ISI)^49^. For each animal, we computed inter-spike intervals for each neuron and fitted the ISI distribution with an exponential function. In response to FAMS, ISI distributions exhibited a distinct exponential decay in all three animals (**Figure 4E, Figure S2D**), indicating quasi-stochastic firing behavior of recruited neurons (coefficients of determination R^2^ = 0.89, 0.63, 0.90 for Ct-Fi, Ct-Lu, Ct-Re respectively). Quasi-stochastic firing was consistently present at all FAMS parameter combinations that evoked sustained firing (**Figure 4F, Figure S2D**). We conclude that FAMS achieves desynchronized population firing and approximates stochastic single-axon firing, and therefore represents a significant step toward inducing more naturalistic activity with electrical stimulation techniques.

### FAMS generates smooth firing rate profiles that resemble natural activity

With rectangular stimulation pulses, each neuron will either not respond or will fire a single action potential in response to each pulse. As such, the neural firing rate is a direct consequence of the stimulation frequency. Conversely, using waveforms based on continuous sinusoids complicates the relationship between stimulus and neural firing patterns. We therefore asked whether it is possible to control the firing frequency of individual neurons by tuning the FAMS waveform parameters. In the same feline preparation, we generated FAMS waveforms of different amplitudes, while keeping the carrier and beat frequency constant (1000 Hz and 10 Hz respectively), and characterized the evoked neural activity. Firing rate profiles exhibited a smooth increase and decrease that was coherent with the beat frequency of the stimulation waveform (**Figure 5A**, (ii)). For each neuron, an increase in stimulus amplitude caused an earlier firing onset and higher firing rates during the whole duration of the stimulation waveform. At the population level, higher current amplitude had two main results: an expansion in the number of recruited fibers (**Figure S4A**), and an increase in the average firing rate over the entire duration of the stimulation waveform (**Figure 5A**, (iii)). Each neuron exhibited a specific relationship between current amplitude and firing rate. We found that most neurons showed a monotonic increase in firing rate with increasing current amplitude (**Figure 5B,C**). Moderate increases in current amplitude induced firing rates lower than 100Hz which is within the range of firing rates that are common during sustained natural firing. Conversely, higher currents generated firing rates higher than 100 Hz, that are typical of transient and rapidly adapting firing^50^. Moreover, in each animal, we found three classes of neurons that exhibited either weak, moderate, or steep increases in firing rate with increasing current (**Figure 5B**). The majority of neurons showed either weak or moderate increases, with maximum firing rates not exceeding 150 Hz. A minority of neurons (10%, 17% and 8% in Ct-Fi, Ct-Lu and Ct-Re, respectively) reached higher firing rates. We compared firing patterns with other state-of-the-art biomimetic waveforms, such as high-frequency and amplitude-modulated rectangular pulse sequences (BioS) ^26^. We found that while BioS could introduce a certain degree of desynchronization (**Figure S3D**), it did not allow us to easily modulate the firing rate of most recruited neurons (**Figure S3 C,E,F**). Indeed, the quick subsequent rectangular pulses in BioS elicited very high firing rates (∼ 200 Hz, **Figure S3 C,E,F**), constraining the neurons to generate a spike as soon as the refractory period was over. Since sustained firing rates in intact neural populations rarely exceed 100 Hz ^50^, we conclude that FAMS outperforms current state-of-the-art biomimetic stimulation waveforms, as it is able to achieve quasi-stochastic and asynchronous firing while also evoking physiological and easily controllable firing rate profiles in more than 80% of the recruited neural population.

**Figure 5.**
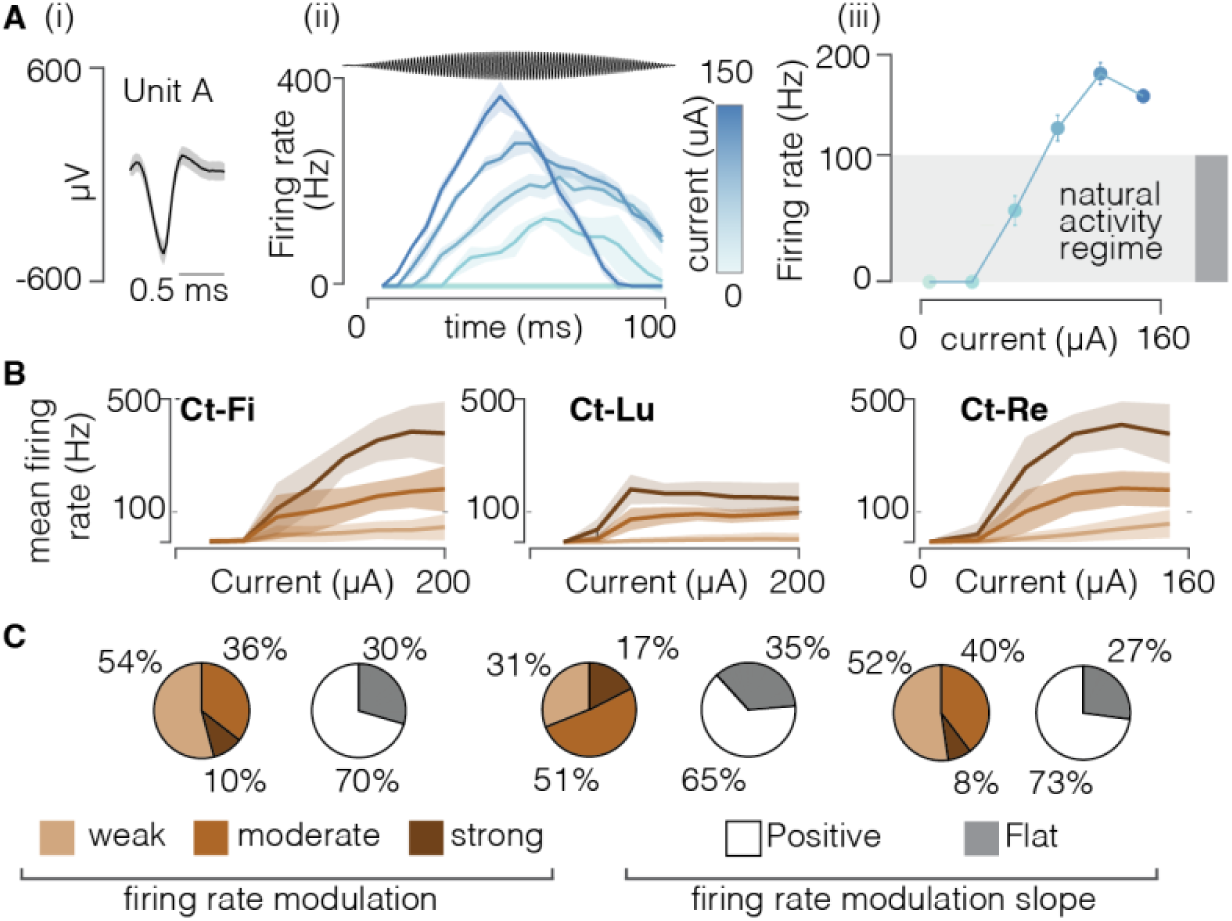
FAMS generates controllable and smooth firing rate profiles. **(A)** (i) Voltage profile of a selected single unit. Solid black line is the mean voltage profile, shaded area represents the SEM. (ii) Firing profiles over time of the selected single unit during FAMS stimulation, color coded as a function of peak current amplitude. Higher currents elicit earlier firing onset and higher firing rates during the whole duration of the stimulation waveform. Solid line is the mean profile, shaded area represents the SEM at each time point. The gray area shows the physiological range for sustained firing rate in mammal (from Wellnitz 2010) (iii) Mean firing rate, averaged across the dura tion of the waveform, for the selected unit, as a function of stimulus amplitude. Increasing the stimulus amplitude evokes higher firing rates, before saturating around 200 Hz. Dots are mean values, bars are SEM. **(B)** Firing behavior during FAMS at the population level. We used firing rate profiles as a function of current amplitude for each unit to group units in three clusters, color coded. Solid line is the firing rate as a function of current amplitude, averaged across all units belonging to the same cluster. The shaded area represents the SEM at each current value. **(C)** For each animal, pie charts representing the assignments of all units to a specific cluster, alongside a second pie chart representing the fraction of units that displayed a positive or flat (weak) correlation of average firing rate with increasing current amplitude. No neurons displayed a negative correlation between average firing rate and current amplitude.

### Amplitude, carrier frequency, and beat frequency of FAMS control neural firing

The FAMS waveform can be tuned according to three main parameters: the peak amplitude, the carrier frequency, and the beat frequency. We explored the influence of each parameter on the firing rate, latency, and desynchronization of the evoked neural activity. We first analyzed the effect of parameter variations on the firing profiles of individual neurons (**Figure 6A** (i)). Increasing the peak amplitude of the FAMS waveform simultaneously increased the average firing rate throughout the duration of the waveform and decreased the response latency (**Figure 6A** (i), **Figure S5B**). This result is easily interpretable, since a higher peak current amplitude is associated with a faster increase in current during the FAMS waveform. Conversely, higher carrier frequencies generated lower firing rates overall (**Figure 6A** (ii), **Figure S5B**). The opposing effects of peak current amplitude and carrier frequency on the evoked firing rates are an important advantage of this stimulation waveform and could be exploited to exert fine control over the whole population. Indeed, while increasing peak current amplitude has the effect of increasing firing rate, it also increases the number of recruited fibers (**Figure S4A**). Decreasing carrier frequency also generates a similar average firing rate increase. Moreover, response timing can be controlled by modulating the beat frequency parameter (**Figure 6A**, (iii)). Indeed, the beat frequency determines the overall duration of the stimulation pulse, and consequently the duration of the evoked neural response. Interestingly, beat frequency also impacted the average firing rate, with higher beat frequencies associated with shorter but weaker neural responses (**Figure 6A**, (iii)). Therefore, for a higher beat frequency, a higher peak current is necessary to achieve short spiking bursts at firing rates that are comparable to a longer and slower FAMS waveform. We also verified that these results hold for the entire population of recruited neurons. Indeed, population activity rose with increasing peak current and dropped with increasing carrier and beat frequencies (**Figure 6B, Figure S5A**). Overall population desynchronization never dropped below an average value of 3.5 ms (Ct-Fi = 5.1± 4.3 ms, Ct-Lu = 3.5±1.9 ms, Ct-Re = 3.9±3.7 ms), with higher desynchronization values associated with lower firing rates (**Figure 6C, Figure S2C, Figure S5B**).

**Figure 6.**
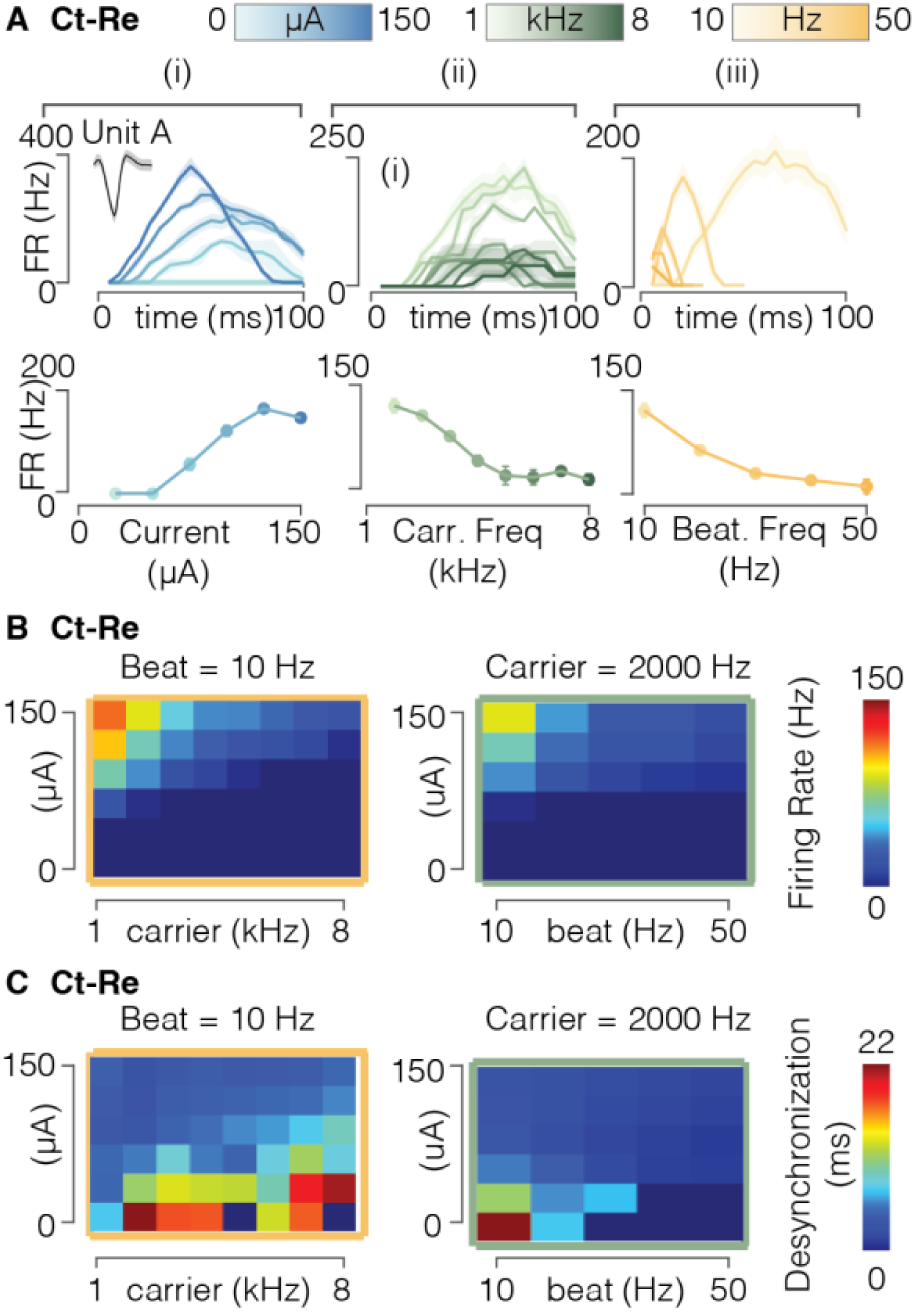
FAMS parameters allow for fine tuning of neural firing. **(A)** Top row: For the same unit selected in Figure 5 (Unit A), firing rate profiles over time, averaged across stimulus repetitions, of the selected single unit during FAMS stimulation, color coded as a function of (i) stimulus amplitude, (ii) carrier frequency, and (iii) beat frequency. Bottom row: For the same selected unit, mean firing rate, averaged across repetitions and the duration of the waveform, for the selected unit, as a function of (i) stimulus amplitude, (ii) carrier frequency, and (iii) beat frequency. **(B)** Heatmap of mean firing rate values for all current amplitude and carrier frequency combinations, as well as all current amplitude and beat frequencies combinations. **(C)** Heatmap of mean desynchronization for all current amplitude and carrier frequency combinations, as well as all current amplitude and beat frequencies combinations.

### FAMS produces more naturalistic percepts when compared with intensity matched biphasic square pulses

We next tested whether the firing pattern differences observed in our in-silico and in-vivo experiments could influence evoked perceptions in a non-invasive peripheral nerve stimulation experiment. To assess the quality of the sensations evoked by FAMS in human participants (**Figure 7A**), we stimulated afferent fibers of the median nerve in the wrist and asked participants (N=10) to evaluate the naturalness of the resulting sensations using a two-alternative forced-choice (2AFC) task (**Figure 7B**). We directly compared two intensity-matched waveforms: (1) FAMS with 2 kHz carrier and 20 Hz beat frequency, (2) biphasic square pulse with a 250 µs per phase pulse duration and repetition frequency of 20 Hz. We first found threshold currents for each waveform, defined as the minimum current required to elicit any sensation in the hand. Threshold measurements were repeated at the end of the experimental procedure, confirming robust stability of the sensory threshold (**Figure 7B**, bottom right panel). Participants were then asked to indicate where they perceived the evoked sensation for each stimulation paradigm. **Figure 7C** shows that the evoked percepts were localized to approximately the same hand area for each participant across the two waveforms. To assess any differences that might occur as stimulation intensity increased, we then used three supra-threshold stimulation amplitudes (1.1×, 1.2× and 1.3×) for each waveform. As a result, we only compared stimulation waveforms that were intensity-matched (i.e., they felt equally intense). Participants performed a 2AFC task where they reported which waveform was more natural. The delivery of stimulus waveforms was randomized between the two stimulus intervals. At each stimulation amplitude, the FAMS waveform was rated as more natural than a rectangular-pulse waveform (**Figure 7D**, p<0.0403, 1.2× p< 0.0079, 1.3× p<0.0008). Moreover, the group-averaged ratio of naturalness rating for FAMS over biphasic rectangular pulses increased as stimulation intensity increased. At the highest stimulation intensity tested, participants choose FAMS as feeling more natural than biphasic square pulses 79% of the time, on average. In summary, human participants consistently rated FAMS-evoked sensations as more natural than sensations evoked by rectangular pulses.

**Figure 7.**
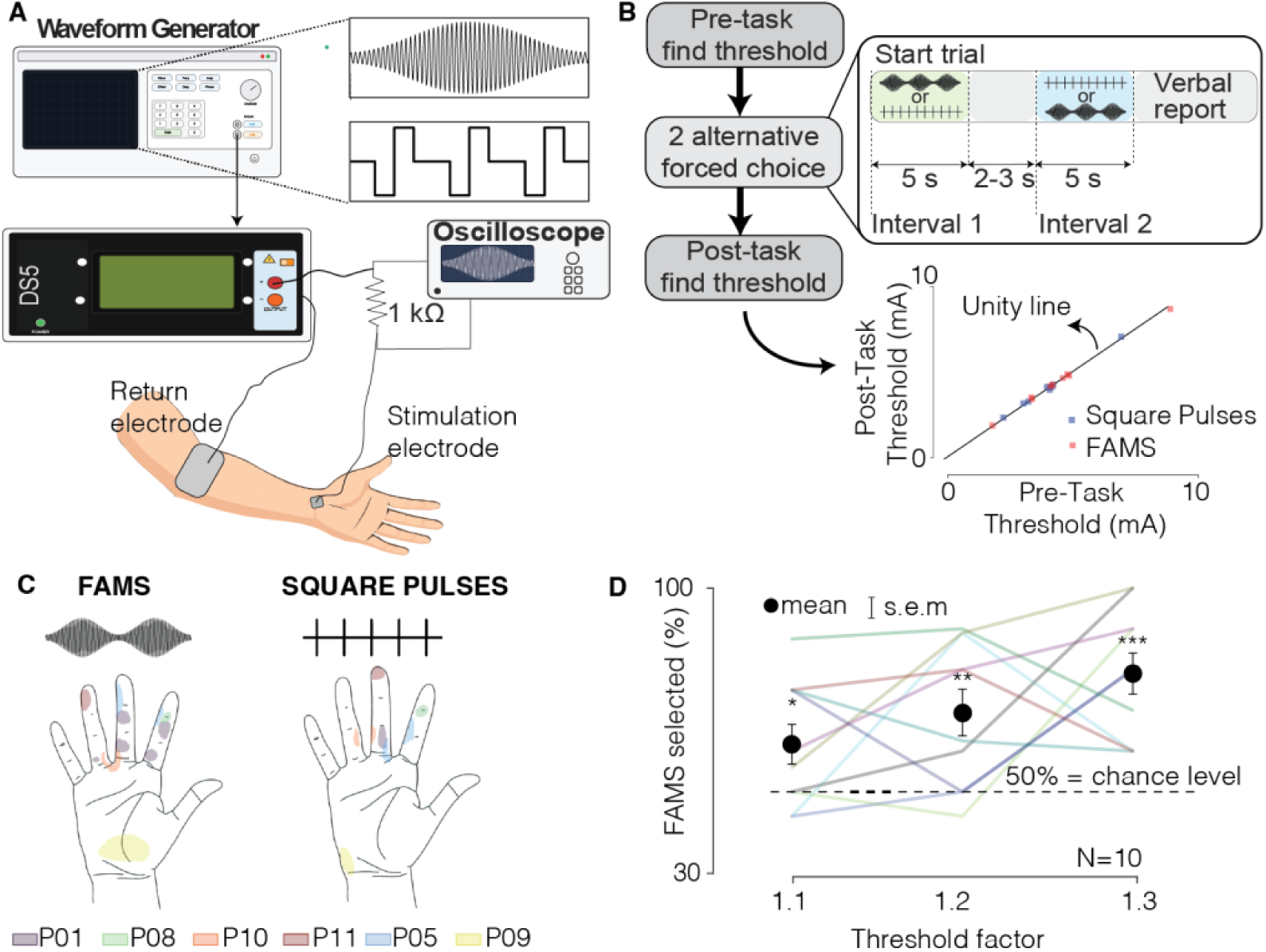
Participants consistently rate FAMS as more natural than intensity-matched rectangular biphasic pulses for median nerve stimulation. (**A)** Experimental set up:. A FAMS waveform or train of bi-phasic rectangular pulses was created using a waveform generator, to control the output of a constant-current source. Stimulation was delivered transcutaneously at the participant’s wrist to stimulate afferent fibers of the median nerve. Both waveforms had a 20 Hz repetition rate. The electrode placed at the wrist stimulated afferent median nerve fibers, and the return electrode’s increased surface ensured no stimulation of the bicep occurred. The output current of the stimulator was measured as the voltage drop across a 1kOhm resistor using an oscilloscope to confirm waveform shape and amplitude. (**B**) Schematic of the experimental procedure. First, we found the sensory threshold for each stimulation waveform. Then, we asked participants to perform a two-alternate forced choice task (2AFC), where they were presented with two intensity-matched stimuli and were asked which felt more natural. Third, we repeated the sensory threshold procedure; The bottom panel shows the relationship between pre-task and post-task measured thresholds. (**C**) Map of sensory percepts. Each colored areas represents the sensory percept of each subject. (**D)** Group-averaged preference for FAMS over biphasic rectangular pulses (percentage of trials; black squares) at three suprathreshold intensities (1.1×, 1.2×, 1.3× of individual thresholds); error bars represent standard error of the mean (N=10 participants; 10 trials per intensity per participant). The horizontal black dashed line indicates 50% chance of choosing FAMS or biphasic rectangular pulses. Also plotted, data for each participant averaged across all ten trials at each threshold factor.

## DISCUSSION

In this study we presented a novel electrical stimulation waveform that evokes irregular and asynchronous yet controllable neural firing. Electrical stimulation is arising as a promising clinical solution for a variety of disorders, such as spinal cord injury^1,6,7^, stroke^8,9^, or limb amputation^10,14,16,17,27^. While stimulation paradigms have proven extremely effective at partially restoring sensory-motor function in animal models and humans, positive outcomes are typically accompanied by some negative effects. These negative effects include production of unnatural paresthetic sensory percepts in people with limb amputation and rapid muscle fatigue during functional electrical stimulation, which are both potentially linked to the highly synchronous firing patterns evoked by traditional rectangular pulses^10,28,32,33^. In particular, recent literature has highlighted that asynchronous firing could decrease paresthetic percepts^28^. Therefore, the development of stimulation waveforms that evoke more physiological firing patterns could constitute a powerful tool to enhance and refine electrical stimulation therapies.

In this study, we found that amplitude modulated high-frequency sinusoidal waveforms can elicit less synchronous firing patterns in neuronal populations. We showed that sinusoidal waveforms are very effective at desynchronizing neural activity. We demonstrated that ion channel dynamics rectify and low-pass filter sinusoidal stimulation profiles, allowing neurons at different locations and with different geometric properties (e.g., axon diameter) to fire at different times. For this reason, lower frequencies are very effective at desynchronizing spiking across populations of neurons. Conversely, higher frequencies are more effective at inducing less regular firing within each neuron. We found that a high-frequency sinusoid, modulated in amplitude at a lower sinusoidal frequency combines those two properties, and generates irregular firing for each neuron, while also desynchronizing spikes at the population level. Importantly, we showed that while the precise time of spikes is quasi-stochastic, the overall firing rate of each neuron and of the whole population is easily controllable by modulating stimulation parameters, namely current amplitude, carrier frequency, and beat frequency. In particular, an increase in current amplitude yields a higher average firing rate. A reduction in carrier frequency also causes an increase in the average firing rate, while also slightly reducing the overall desynchronization. During FAMS, firing profiles exhibited both a gradual rise and descent phase, and the steepness of those phases depended on the beat frequency. The ability to control all aspects of firing rate profiles with only three parameters allows for simple stimulation designs, which are more likely to succeed in clinical settings. Moreover, as our biophysically realistic computational model accurately captures the firing dynamics of a neuron population with variable diameters (**Figure S6A,B**) we argue that a computational model could be a valuable tool to efficiently design stimulation protocols based on the FAMS waveform to achieve specific firing patterns targeted to specific therapeutic applications.

Extending our exploration to human subjects, we found that participants rated FAMS as being more natural compared to biphasic pulsed current stimulation, across a range of supra-threshold stimulation amplitudes. We suggest that the increased naturalness rating observed for FAMS waveforms over traditional biphasic square waveforms originates from differences in the evoked firing patterns in response to FAMS. The less synchronized firing patterns of the FAMS waveform may better approximate naturalistic firing patterns and, as such, are perceived to be more naturalistic. These data suggest that adoption of FAMS may reduce some of the unwanted and uncomfortable sensations which can be induced by standard biphasic square pulses, while also allowing for careful control of firing patterns using the various waveform parameters which define the FAMS waveform (e.g. amplitude, beat frequency, and carrier frequency).

Several important limitations remain to be addressed in future studies. First, the safety of using waveforms based on sinusoidal profiles, especially with implanted electrodes in close proximity to neural tissues, requires further investigation. Studies that have characterized electrode and tissue damage associated with the repeated delivery of rectangular pulses^24^ would need to be replicated for the use of sinusoidal waveforms. Similarly, the existing safety limitations on the usable parameter space would need to be reevaluated. This is pivotal to guarantee that neurons are not driven to activity levels that lead to aberrant axon and network activity^51^. Second, a paradigm shift from rectangular pulses to sinusoidal profile would require appropriate stimulation technologies. Most implantable stimulators are currently designed to deliver pre-programmed rectangular pulses, whose amplitude and frequency can be controlled in real time with an external control signal. Analog sinusoidal waveforms, such as FAMS, would require delivering continuous waveforms with a fine temporal resolution. The growing number of studies employing sinusoidal stimulation waveforms in various forms and applications^52–56^ underscores a field-wide demand for stimulators capable of safely and efficiently delivering continuous, high-fidelity analog waveforms, suggesting that the development of such technology is both necessary and forthcoming. Finally, the performance of FAMS relative to more traditional stimulation waveforms was assessed only in the context of transcutaneous stimulation of the median nerve in the arm. Although these findings suggest that FAMS may be a promising strategy for generating biomimetic neural activation, this waveform must be carefully evaluated across a broader range of sensory and motor applications. In particular, the effects of FAMS on evoked sensory percepts when stimulating different regions of the nervous system, its influence on evoked paresthesia, and its impact on the modulation of muscle contractions should be thoroughly characterized to ensure optimal clinical outcomes.

In the past few years, other studies have proposed alternative stimulation waveforms to achieve asynchronous and stochastic neural firing. Those methods pioneered the notion that, by gradually increasing^26^ and/or decreasing^40^ the amplitude of pulses in a high-frequency train, neurons would be gradually recruited by each pulse based on their specific physiological and geometrical properties. While modulating the amplitude of rectangular pulses did indeed introduce a degree of desynchronization in the resulting neural activity, the use of rectangular pulses limits the degree of desynchronization and often generates firing rates that are not controllable and are above the physiological range for sustained firing. In fact, desynchronization through the use of square pulses is achieved by overstimulating fibers during their refractory period^28^. Conversely, FAMS achieves desynchronization by using sinusoidal current delivery while also allowing the necessary control over firing rates to encode information as required by each specific application. In conclusion, FAMS represents an important advancement in the design of biomimetic stimulation waveforms to evoke physiological firing patterns which are easy to control and implement. The FAMS waveform could notably improve the outcome for clinical applications in which the naturalness of evoked neural activity is of particular importance, such as restoration of touch and proprioceptive sensations in people with sensory deficits.

## MATERIALS AND METHODS

### Computational model

#### Electrical stimulation

To model the extracellular potential fields generated by the stimulation modalities examined in this study, we considered one bipolar electrode, with contacts located at [5, 1, 0] mm from the origin. We modeled the electrodes as ideal point sources and we calculated the potential field under quasistatic conditions (Plonsey and Heppner, 1967). A more detailed description of the implementation of this model is available in our previous work^44^. We simulated a variety of different waveforms: single, symmetric, biphasic rectangular pulses (pulsewidth 50 µS, and variable amplitude); sinusoidal waveforms (variable amplitudes and frequencies = 50, 100, 500, 1,000 Hz); FAMS waveform, which is expressed by time-varying equation (1):

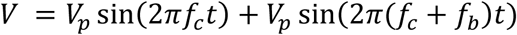

where V_p_ is half of the resulting peak voltage amplitude, f_c_ is the high (*carrier*) frequency, and f_b_ is the low (*beat*) frequency. To characterize and compare neural activations in response to sinusoids of different frequencies, we computed the threshold current amplitude for an axon of diameter equal to 10 μm, extending along the x axis and located at y = 0 and z = 0. We defined the threshold current amplitude as the minimal current amplitude required to generate a single spike. We then used 1.5 times the threshold current amplitude to simulate both single cells and populations responses.

#### Biophysical neuron models

This method has been adapted from Mirzakhalili et al, 2020^44^. We modeled the electrical properties of nerve or white matter tissues as an anisotropic infinite volume. We modified a previously existing compartmental model of a myelinated mammalian axon, that utilized the standard Hodgkin-Huxley formalism ^57^ by adjusting the functional form of the kinetics for the slow potassium channel ^44^ and then applied the spatiotemporal voltages described in the previous section. We used the morphology of the original compartmental model of a myelinated mammalian axon (McIntyre et al., 2002). This morphology models different sections of the myelin along the internode by including myelin attachment segment (MYSA), paranode main segment (FLUT), and internode segment (STIN) regions of the fiber. We simulated neurons with diameters comprised between 6 and 12 µm. To calculate the neural response to electrical stimulation we solved the time-dependent partial differential cable equations using the NEURON software package ^46^. A more detailed description of the implementation of this model is available in our previous work ^44^, (STAR Methods, sections, *Myelinated Axon Model, Morphology, Ionic Currents, Model Implementation*).

#### Analysis of irregular and regular firing in single axons

To automatically compute the duration of irregular firing during electrical stimulation, we calculated a sample cross-correlation ^58^ in MATLAB (Mathworks) by sliding a single spike template over the entire membrane voltage trace. We chose one spike during regular firing as the template. We then identified all local maxima of the cross-correlation index to look at the overall evolution of the similarity of the spikes to the template over time. We computed the time of regular spiking onset as the first time instant at which the cross-correlation index constantly reached the value of 1.

#### Spectral analysis of membrane currents

To quantify the amount of stimulus rectification performed by the neuronal membrane, we computed the fast Fourier transform (FFT) of the transmembrane current in MATLAB (Mathworks). The frequency content of the transmembrane current time profiles reveals how the axonal membrane filters and demodulates the stimulus ^44^. For example, as we deliver a stimulus that only includes high-frequency content (at 1000Hz and 1010Hz), the amount of demodulation performed by the membrane would be indicated by the amplitude of the power spectrum at 10 Hz (i.e. the difference between the 2 frequency components). We therefore extracted the FFT amplitude value at 10 Hz and to quantify of the amount of demodulation that the neuronal membrane generated during FAMS stimulation.

### Surgical procedures

We conducted acute experiments in three adult male cats. All procedures were approved by the University of Pittsburgh Institutional Animal Care and Use Committee and were performed according to all guidelines and regulations. We induced anesthesia with a combination of ketamine and acepromazine and maintained a surgical anesthetic plane with isoflurane (1-2%). We performed a tracheotomy and connected the trachea to an artificial respiration system that ventilated the animal at 12-14 breaths per minute during the duration of the procedure. Temperature was maintained using a warm air heating pad while IV fluids were continuously administered. We used a catheter in the carotid artery to monitor blood pressure continuously throughout the experiment. At the end of the experimental procedure, animals were euthanized through an IV injection of Euthasol. We performed a laminectomy to expose the L6 and L7 dorsal root ganglia. Using a high-speed pneumatic inserter, we implanted 32-channel multi-microelectrode arrays (Blackrock Microsystems) in the DRG at the L6 and L7 segments. Each electrode shank was 1 mm long with a 50 µm exposed tip and inter-electrode pitch of 400 µm. The arrays implantation was achieved using a pneumatic compressor system (Impactor System, Blackrock Microsystems). A stainless-steel wire with ∼1 cm of insulation removed served as a ground electrode and was placed subcutaneously in the lower back. To perform electrical stimulation, we placed a multi-contact stimulating cuff electrode (Ardiem Medical, Indiana, PA) around the sciatic nerve. The cuff electrode was comprised of 12 contacts, divided in three rings of four contacts each. Rings were 4 mm apart in the direction longitudinal to the nerve, and contacts within each ring were 1.2 mm apart (**Extended Data Figure 2B**).

### Electrical stimulation of the sciatic nerve

We delivered stimulation through an analog stimulus isolator (Model 2200, AM systems, Sequim, WA), controlled by a custom voltage signal. The custom control signal was digitally designed and then generated using multifunction input/output device (Model USB 6356, National Instruments, Emerson, USA). All waveforms were preprogrammed through a custom Python application, and then transferred to the I/O device that controlled the analog stimulus isolator in real time. Applied currents ranged from 10 to 400 µA, and frequencies ranged from 1 to 8000 Hz. In all stimulation conditions, the inter-pulse interval was equal to 1 s. We delivered stimulation from two contacts, longitudinally disposed and aligned along the sciatic nerve, 8 mm apart (**Extended Data Figure 2B**). For all animals, stimuli were delivered in the same order. First, we delivered rectangular pulses at 1Hz and different amplitudes to qualitatively identify the threshold and saturation current amplitude. We then delivered FAMS stimuli of increasing amplitude, increasing carrier frequency and increasing beat frequency.

### Manual stimulation of the sciatic nerve

In order to observe the baseline activity of the recorded units, we performed electrophysiological recordings while the leg of the animal was repeatedly flexed and extended by handling the foot of the animal and rotating the lower leg around the knee. We repeated these movements on average every 3-4 seconds, for seven consecutive times, at the beginning of the experimental session. This procedure was performed for animals Ct-Lu and Ct-Re.

### Electrophysiological recordings from the dorsal root ganglia

We recorded multiunit activity from each electrode tip, for a total of 64 voltage traces per animal. We digitized and sampled neural activity at 30 kHz with a Nano2 Headstage (Ripple LLC) and a Grapevine Neural Interface Processor. Signals were then filtered with a 3^rd^ order high-pass Butterworth filter, with a 0.1 Hz cutoff and a low-pass filter with cutoff at 7,500 Hz.

### Human Participant Experiments

#### Ethics

Experiments on healthy volunteers were performed in accordance with approval from the CEITEC Ethical committee for research on human participants. Stimulation was delivered using the Digitimer DS5 device which is clinically approved in the EU.

#### Participants

Participants were recruited via advertisements on the local campus as well as through an email list for people who had previously expressed interest in prior studies. A total of 13 participants were recruited for these experiments with a final sample size of N=10 being included in the experiment (30.4 ± 11.6 years; mean age ± standard deviation, 40% Female). In general, the non-dominant hand of the participant was stimulated, except in cases where there was a history of sports injuries in the wrist or nerve injuries involving the carpal tunnel. Participants were not compensated for their participation. Each participant gave informed written consent. Participants were excluded for one of two reasons; for one participant they requested to stop the experiment during the threshold finding procedure, and the other two participants were excluded due to excessive movements and repositioning of the arm affecting the established threshold values. These participants did not have enough time to be remeasured, so they were excluded from the study.

#### Electrical stimulation hardware

Simulation was delivered using the Digitimer DS5 constant current source. Gel-assisted carbon transcutaneous electrical nerve stimulation (TENS) electrodes were used for stimulation. For the active electrode, TENS electrodes were cut to a small size (approx. 3 cm^2^) to fit over the carpal tunnel. A much larger TENS electrode (10×5 cm) was used as the return electrode, placed on the bicep. Stimulation waveforms were produced with a two-channel Keysight EDU33212A function generator and delivered through a bipolar constant current stimulator (DS5) to provide constant-current output. Because the DS5 exhibits frequency-dependent attenuation above 1 kHz (per the device documentation), we did not rely on the set value and instead verified the delivered current by measurement Output current was always measured as the voltage drop across a calibrated 1 kΩ resistor, using a digital oscilloscope (Picoscope 2206D, input impedance 1 MΩ, software: PicoScope 7 v7.1.39.3737). A schematic representation of the stimulation circuit can be seen in figure 7A. All currents reported as part of the human stimulation experiments are these measured values in mA. The stimulation experiments were controlled using Python code via PsychoPy to control experimental parameters and waveform generator. This allowed experimental parameters to be randomized.

#### Experimental procedure

Stimulation targeted afferent sensory fibers in the median nerve near the carpal tunnel (**Figure 7A**). Stimulation at this site typically evoked sensations in the fingers and palm of the participant’s hand, with modalities including ‘tingling’ or ‘pins and needles’. The active electrode was positioned above the carpal tunnel along the center line of the arm, with the distal wrist crease serving as a landmark. One edge of the electrode was placed along the distal crease, so the remainder of the electrode sat further from the palm. A second return electrode with a much larger surface area was placed on top of the bicep. This much larger surface area ensured only stimulation of the median nerve fibers at the wrist contributed to the evoked hand sensations.

Two stimulation waveforms were compared: (1) a biphasic square pulse with a cathodic pulse width of 250 μs and a 20 Hz repetition frequency, and (2) FAMS amplitude modulated kHz sine wave with a carrier frequency of 2 kHz (250 μs cathodic phase duration) and a beat frequency of 20 Hz (figure7A).

Participants were seated in a comfortable position with their arm resting on a lightly cushioned surface, such that minimal movement of the arm occurred once the experimental procedure began. A benefit of this stimulation site is that it is relatively robust to small movements in the arm of the participant. However, participants were informed that they should maintain a constant arm position to minimize changes in the effects of stimulation.

To determine the threshold for stimulation for each waveform, an adaptive staircase method (3 up/1 down), with a minimum of ten reversals was used. The step size used was 0.05mA. To determine an approximate threshold to begin the staircase procedure, stimulation amplitude was increased manually until the participants noted they could perceive the evoked sensations, the amplitude was then decreased until the sensations disappeared. At this point the stimulation was stopped, and the staircase method began at this amplitude. We wanted to find the minimum current required to evoke a sensation in the participant’s hand. The median nerve branches quite significantly after passing through the carpal tunnel to innervate large parts of the palm and fingers. Most notably for our experiment, these include the palmar and digital branches, which innervate much of the palm and the fingertips, respectively. To ensure that the same fibers were being stimulated between waveforms, participants were asked to circle on a cartoon drawing of a hand where they felt the stimulation. It was ensured between waveform types that the general area first stimulated was also stimulated with the second waveform, ensuring as fair a comparison as possible (**Figure 7C).** We randomized the order in which we determined the threshold for the two waveforms. Based on the measured thresholds the subsequent intensities (1.1×, 1.2×, 1.3×) were calculated and presented to the participants during the experiment.

A two-alternative forced-choice (2AFC) design, with option to abstain, was employed in this experiment. The option for abstention was only used in the case that a participant did not feel either of the waveforms. Otherwise, a forced choice between the two waveforms was required. Participants were presented with two intervals of stimulation, one containing a train of biphasic square pulses and one containing FAMS, with a 2-3 s pause in between the intervals. During each interval, stimulation ramped up to the desired intensity over 2-3 seconds and then remained constant for 5 seconds before switching off. Immediately following the second interval, the subject reported which interval felt more natural. We tested three different intensities of stimulation: 1.1x, 1.2x and 1.3x above the measured minimal perceptual threshold. Both the order of presentation and intensity level across trials were randomized using a Python (version 3.13) script run in PsychoPy (version 2024.2.4). Each waveform pair and intensity level were presented to each participant ten times. Both participant and experimenter were blinded to the order of presentation and the intensity level. A brief training session (approximately 9 trials) was carried out after thresholds were determined to familiarize the participants with the structure of the experiment. During this time, the experimenter asked the participants: “Are the intensities of the stimulation approximately equal or different?”

At random points, approximately halfway through the experiment and close to the end, stimulation was delivered at threshold for each waveform, and the participants were again asked about the relative intensity of the stimulations. For these specific trials, the experimenter was unblinded as to the waveforms being presented. After the experiment, a shorter threshold-finding procedure, with fewer reversals was used to ensure the threshold levels for each waveform had not significantly shifted during the experiment. In most cases this was a minimum of five reversals, and often much higher, except in two exceptional cases where the number of reversals was fewer than five. Again, participants were asked to draw a circle on a cartoon picture a hand the location of the evoked sensations.

#### Stimulation Hand maps Per Participant

Participants marked the location where they felt the stimulation on a paper print out of the outline of a hand. These were then scanned and traced over, digitizing the images. This was done using custom code developed using Python (version 3.14). This allowed representative shapes capturing the areas highlighted by participants to be overlayed and colored (Figure 7C).

### Data analysis

We applied all neural data analysis techniques offline, with the same procedures used for all three animals.

#### Preprocessing

For each recorded channel, the raw signal recorded during the delivery of different stimulation waveforms and parameters was concatenated into a single voltage trace, and then analyzed. We computed a common average trace, by averaging signals recorded from each electrode. The common average was then subtracted from each channel to eliminate artifactual features of the signal that were common to all recording electrodes. Signals were then filtered with a 3rd order Butterworth digital filter with a 10 – 750 Hz bandwidth in Offline Sorter (Plexon, Dallas, Texas). We also used Offline Sorter to extract single units from the voltage traces. We classified a unit as any waveform crossing a threshold 1.5 times the standard deviation of the whole voltage trace. We manually modified the value of thresholds whenever the predetermined values appeared to be inappropriate to successfully extract single units. Spikes were then sorted into single units using a proprietary clustering algorithm in Offline Sorter. Data were manually reviewed in order to eliminate stimulation artifacts from the sorted units and residual stimulation artifacts were removed from each clusters.

#### Automatic clustering of single units

Units were clustered in distinct groups by analyzing the dependency of their average firing rate on stimulus amplitude. For each unit, we computed the average firing rate at each stimulus amplitude. We then used a K-means algorithm (MATLAB, MathWorks) to cluster units in subgroups. We iteratively repeated this procedure with different number of clusters, ranging from two to ten clusters. For each clustering performed with a specific cluster number, we computed the silhouette measure ^59^ to evaluate the separability of the resulting clusters. We found that three clusters guaranteed the best separability for each animal. We then represented the firing behavior of each cluster separately, as reported in Figure 5.

#### Correlation analysis of single units firing rate

We performed a correlation analysis for each unit to explore the relationship between the firing rate and the current amplitude. We computed the correlation coefficients with MATLAB (MathWorks). If C(x,y) is the covariance of two vectors x and y, we computed the correlation coefficient as:

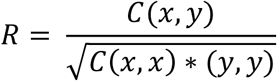

where x is the current amplitude and y is the firing rate of a specific unit. We computed the p-value by transforming the correlation to create a t-statistic having N-2 degrees of freedom, where N is the number of rows of X. We then labeled units as having a *positive* correlation of firing rate with current whenever the correlation coefficient was positive with p<0.05 significance level.

We labeled units as having a *negative* correlation of firing rate with current whenever the correlation coefficient was negative with p<0.05 significance level. All units with positive or negative correlation coefficients with significance levels p > 0.05 were labeled as having a flat dependency.

#### Measure of firing desynchronization

For neural firing evoked by FAMS, desynchronization was evaluated by quantifying the average time distance of each spike from the nearest spike on every other channel. For each spike, generated from each neuron, we computed the time between the selected spike and the closest spike generated by every other neuron. We then averaged this measure to obtain one value associated with each spike. We repeated this procedure for each spike in each dataset. For rectangular-pulse stimulation, we devised a similar metric for the compound action potential response recorded on each DRG microelectrode: we computed the time that intercurred between the beginning of the evoked response for that electrode to the end of the evoked response for every other electrode. We designed this measure to be the most conservative measure of desynchronization for rectangular pulses.

#### Statistics

All data are reported as mean ± s.e.m. or mean ± s.d., and the specific choice is highlighted in the figures or in the relevant caption. If not differently indicated, statistical significance was analyzed using the nonparametric Wilcoxon rank sum test. The level of significance was set at *p < 0.05, **p < 0.01 and ***p < 0.001. In boxplots, the whiskers extend to the most extreme data points, without including considered outliers. An outlier was defined as a point greater than 3 Median Absolute Deviation (MAD) from the median. The MAD was defined as:

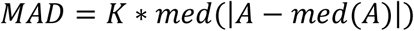

where K is a scaling factor and is equal to approximately 1.48.

#### Human Median Nerve Experiment Statistics

Participant level proportions were calculated for the trials in which FAMS was judged more natural. Normality of these participant-level proportions, at each threshold factor (1.1, 1.2 and 1.3) was assessed, using Q–Q plots (and Anderson–Darling tests). No substantial deviations from normality were observed, therefore, two-sided one-sample t-tests were used to compare mean proportions against chance (0.5) using a p value < 0.05. Because three hypothesis tests were performed (one per threshold factor), Holm correction was applied to control the family-wise error rate at α = 0.05.

## Acknowledgments

We thank Dr. Marco Capogrosso for his help and advice during experimental procedures and the insightful scientific discussions. We thank Rachel Pitzer for help with the animals as well as anesthesia and surgeries setup.

## Author contributions

Conceptualization: BB, LEF

Methodology: BB, RK, CG, LEF

Software: BB, EM

Formal Analysis: BB, DSR

Data Curation: BB, DSR

Writing – Original Draft: BB, LEF, DSR, EDG

Writing – Reviewing & Editing: BB, RK, CG, EM, SFL, RG, LEF, DSR, EDG

Visualization: BB, EM, SFL, LEF, DSR

Supervision: SFL, LEF, EDG

Project Administration: BB, RG, LEF

Funding Acquisition: BB, RG, LEF

## Data and materials availability

All data are available in the main text or the supplementary materials.

