## Supplemental Figures S1-S6 for "High-frequency amplitude-modulated sinusoidal stimulation desynchronizes neural activity and enhances naturalness of evoked sensations"

Barra et al.

The pdf contains Figures S1 to S6.

**Fig. S1.**

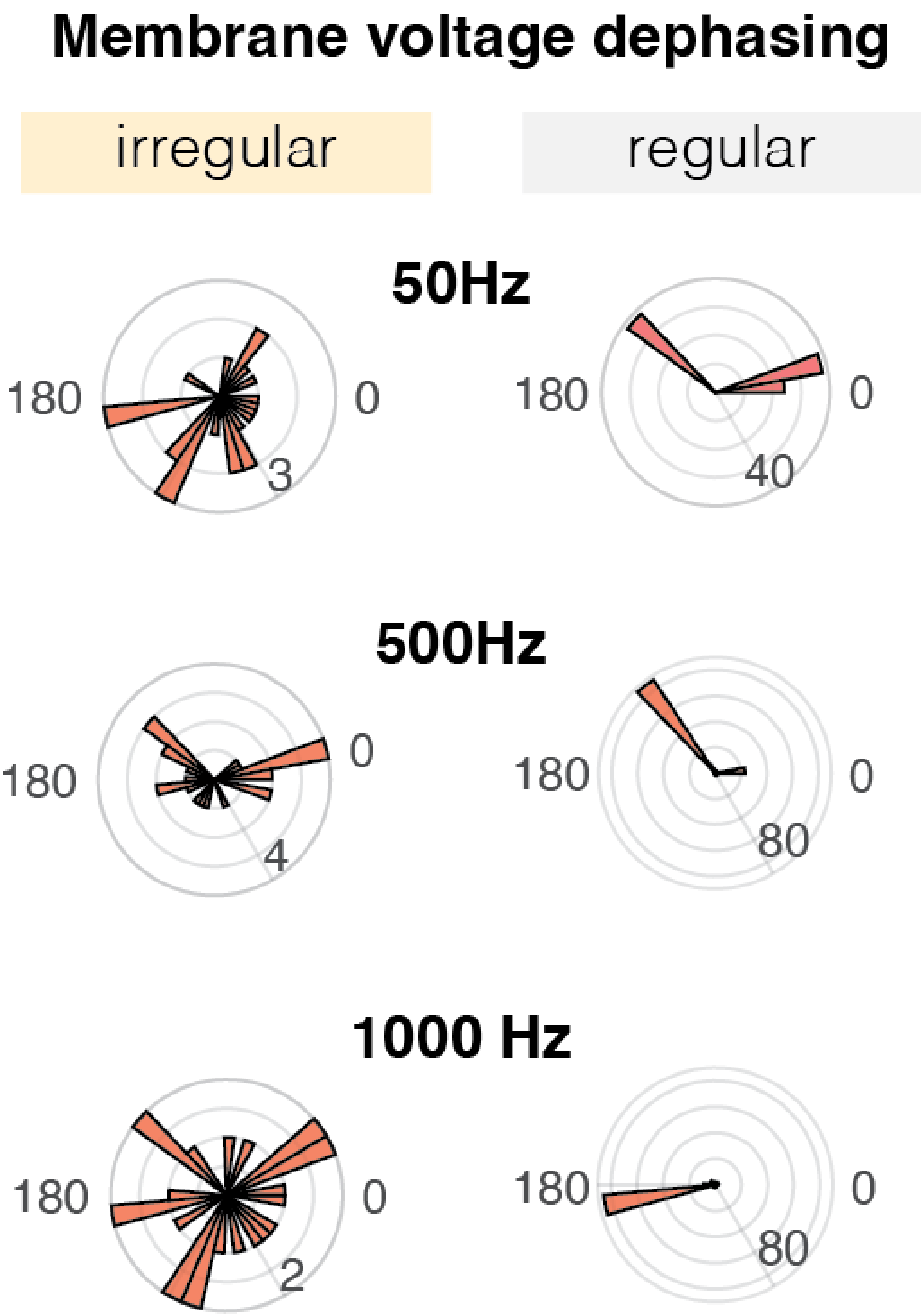


**Figure S1. Sinusoidal stimulation creates phase shifts in membrane voltage oscillations.** Radial histogram of the phase difference between the stimulation waveform and the transmembrane voltage, computed at the time of spike initiation. Left: during irregular firing. Right: during regular firing. The scattered phase distribution during irregular firing indicates that spikes are generated at very different parts of the stimulation waveform.

**Fig. S2.**

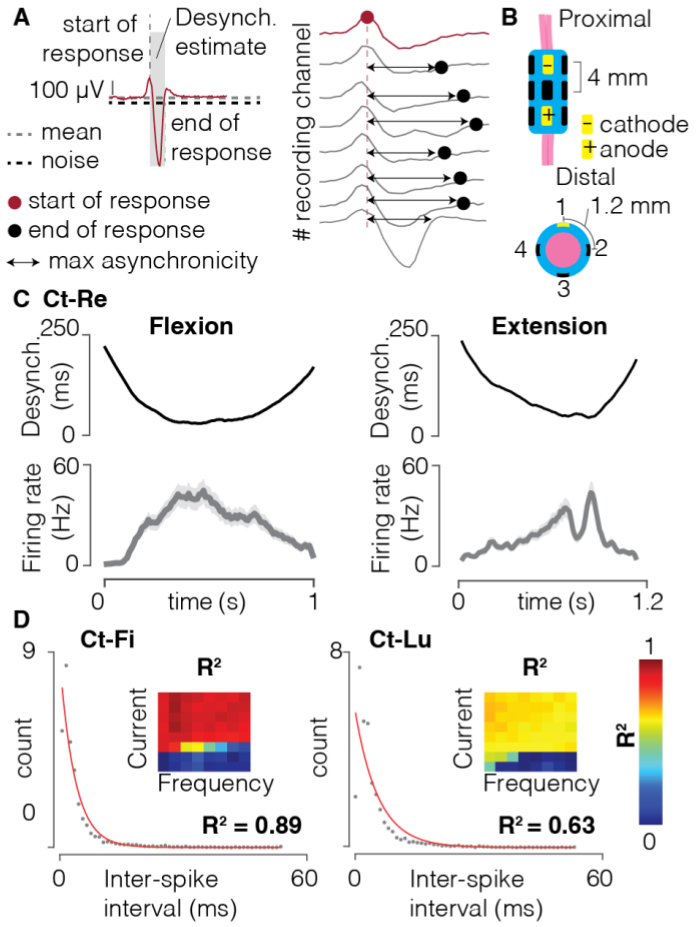


**Figure S2.** Firing desynchronization generated during FAMS and rectangular pulses **(A)** Schematic of desynchronization calculation for rectangular pulse conditions. We used the most generous measure of desynchronization by estimating the time between the pre-hyperpolarization peak recorded on one electrode and the time the voltage returned below the noise level (i.e., 1.5 standard deviations above the mean) on all different electrodes. **(B)** Schematic of the stimulation cuff. The cuff had three rings of four contacts each. The contacts in yellow indicate the contacts used for stimulation, with the cathode closer to the proximal limb and the anode closer to the distal limb. **(C)** Desynchronization and firing rate of a single unit during manual flexion and extension of the animal leg. Solid line: mean trace, averaged across different repetitions of the same movement. Shaded area: standard deviation at each time point, averaged across different repetitions of the same movement. Desynchronization is negatively correlated with firing rate. **(D)** For Ct-Fi and Ct-Lu, scatter plot of the inter-spike interval histogram for Ct-Re (1,000 Hz carrier frequency, 75 µA peak current, 10 Hz beat frequency), along with the relative exponential fit (red line, $R^2 = 0.89$ Ct-Fi and $R^2 = 0.63$ Ct-Lu). Next to each curve, the heatmap of $R^2$ values for exponential fits of ISI histograms at all parameters combination. An exponential distribution of ISIs is indicative of stochastic firing. The $R^2$ matrix indicates that quasi-stochastic firing is obtained for all parameter combinations that elicit sustained firing.

**Fig. S3.**

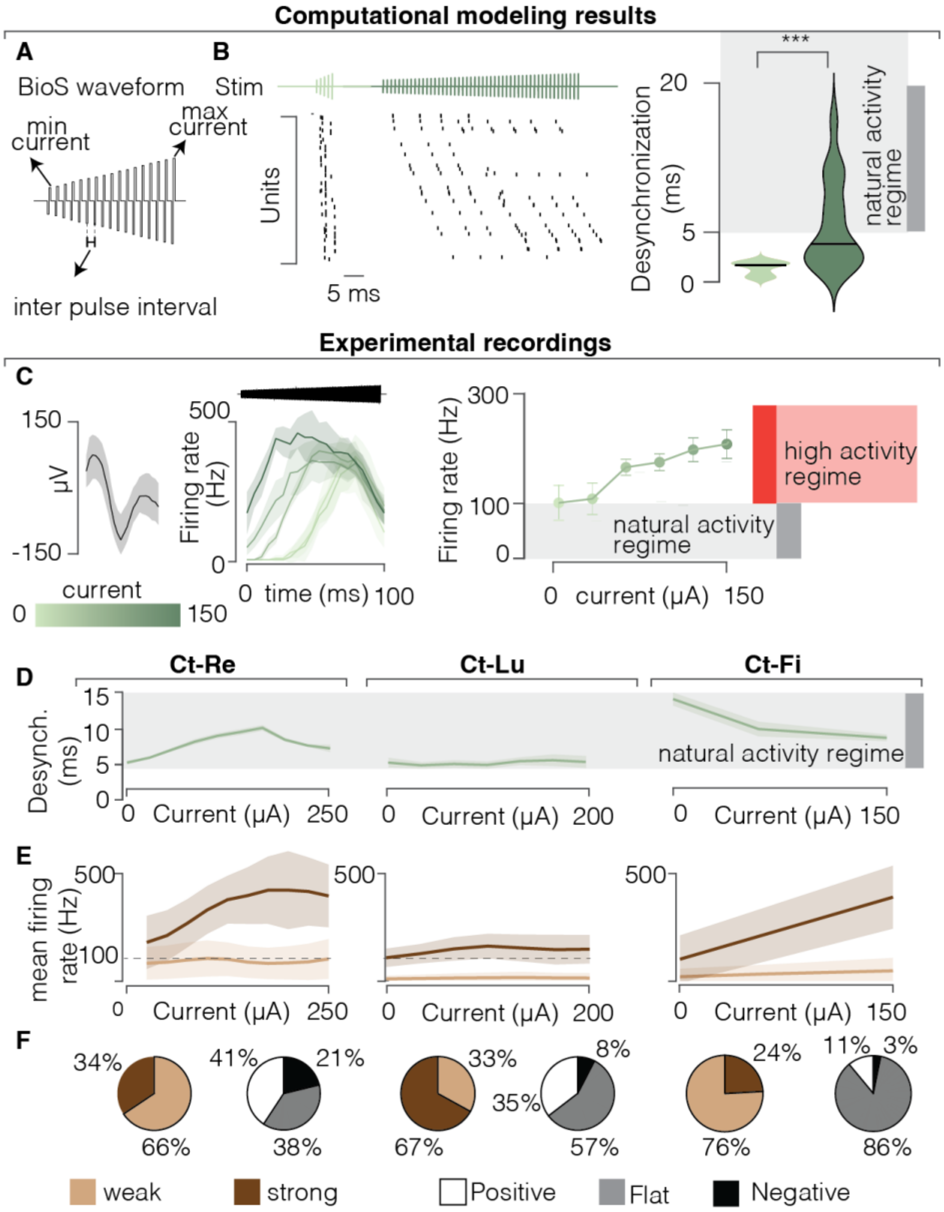


**Figure S3 | BioS generates non controllable and abrupt firing rate profiles | (A)** Illustration of the BioS waveform (Formento, 2020), composed of a series of rectangular pulses of increasing amplitude, delivered at high frequency (>800Hz). **(B)** Raster plot of computationally simulated neurons responding to short and long BioS waveforms, with the same minimum and maximum amplitude. On the right, violin plot of the desynchronization measure for both of these conditions. The shorter waveform achieves less desynchronization, but does not elicit repeated firing in the recruited neurons. The longer waveform evokes more desynchronized firing, but the neurons fire periodically as soon as the refractory period is over, therefore generating unnaturally high firing rates. *** $p = 0.001$, Wilcoxon rank sum test. **(C)** (i) Voltage profile of a selected single unit (same as in Figure 5). Solid black line is the mean voltage profile, shaded area represents the SEM. (ii) Firing profiles over time of the selected single unit during

BioS stimulation, color coded as a function of peak current amplitude. Higher currents elicit earlier firing onset but the firing rates during the whole duration of the stimulation waveform do not change with current amplitude. Solid line is the mean profile, shaded area represents the SEM at each time point. (iii) Mean firing rate, averaged across the duration of the waveform, for the selected unit, as a function of stimulus amplitude. Increasing the stimulus amplitude causes higher firing rates, but even the lowest firing rates are above the normal physiological range of neurons. Dots are mean values, bars are SEM. **(D),** For the three animals, desynchronization as a function of peak current amplitude during BioS stimulation. Solid line is the mean profile, shaded area represents the SEM. **(E)** Firing behavior during BioS at the population level. Firing rate profiles as a function of current amplitude for each unit were used to group units in two clusters, color coded. Solid line is the firing rate as a function of current amplitude, averaged across all units belonging to the same cluster. The shaded area represents the SEM at each current value. **(F)** For each animal, pie charts representing the assignments of all units to a specific cluster, alongside a second pie chart representing the fraction of units that displayed a positive, flat (weak), or negative correlation of average firing rate with increasing current amplitude.

**Fig. S4.**

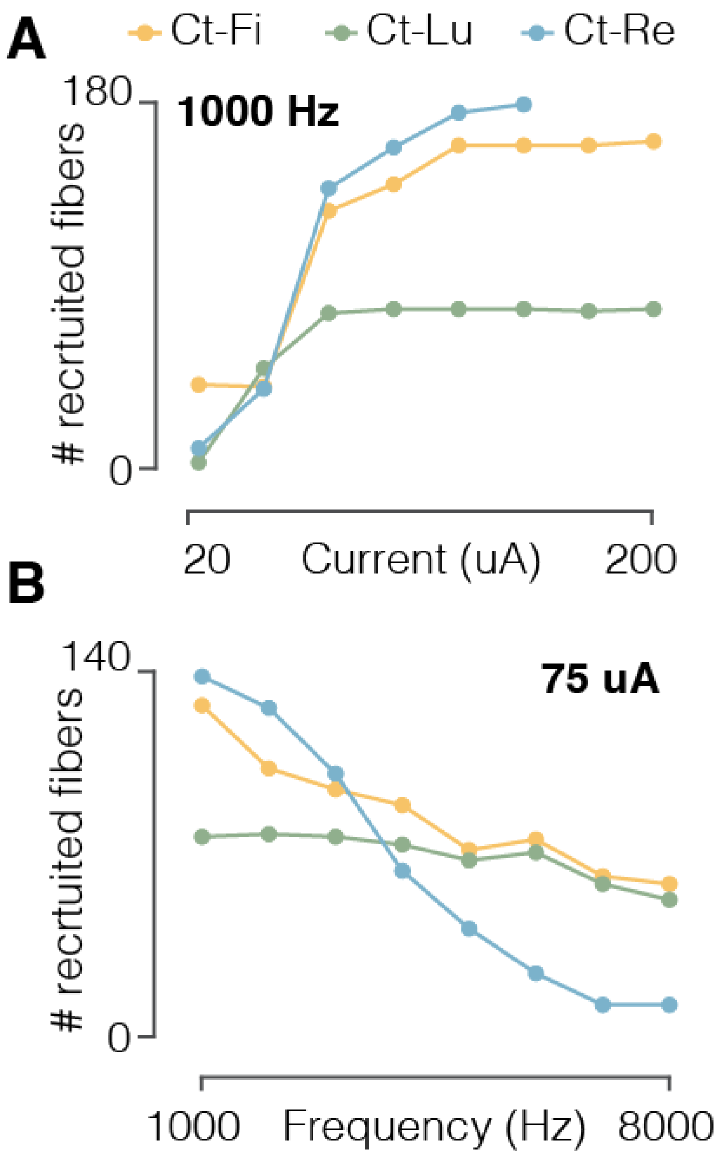


**Figure S4. FAMS recruits more axons at higher stimulus amplitudes (A)** For the three animals, number of recruited fibers as a function of stimulus amplitude. **(B),** For the three animals, number of recruited fibers as a function of stimulus carrier frequency.

**Fig. S5.**

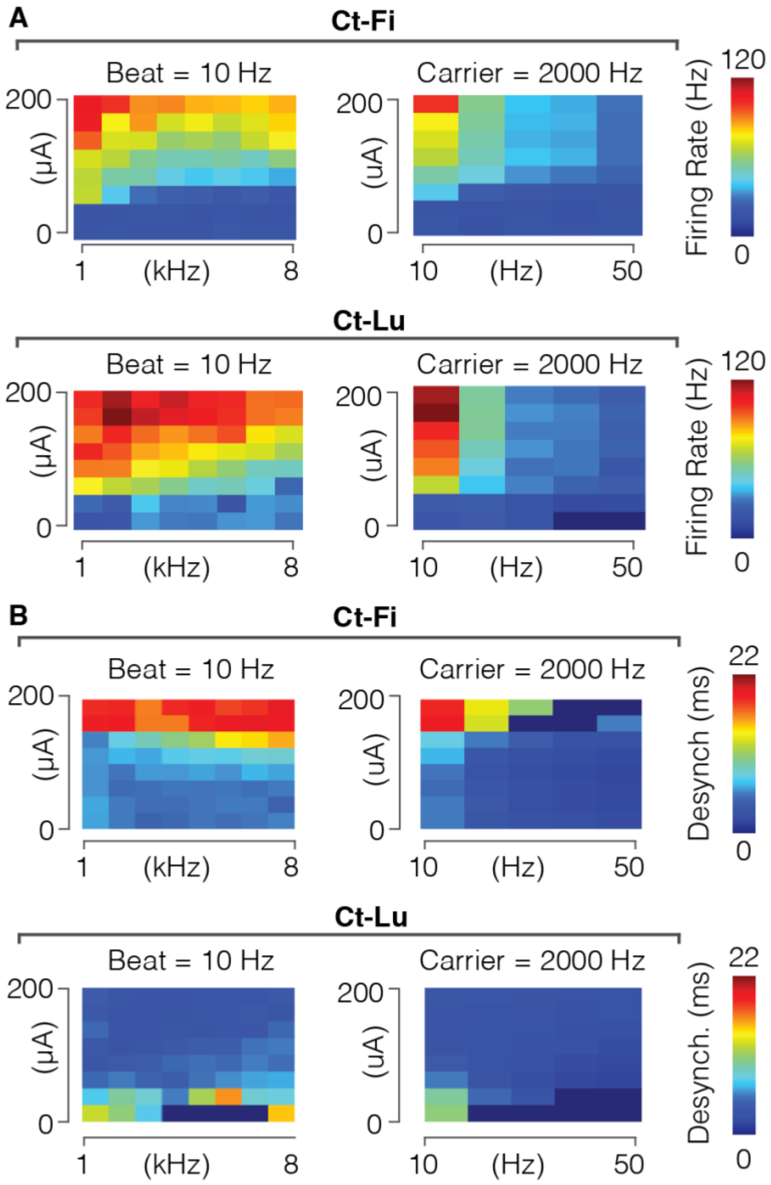


**Figure S5. Firing rate and desynchronization depend on current amplitude, carrier frequency and beat frequency. (A)** Heatmap of mean firing rate values for all current amplitude and carrier frequency combinations, as well as all current amplitude and beat frequencies combinations, for Ct-Fi and Ct-Lu. **(B)** Heatmap of mean desynchronization for all current amplitude and carrier frequency combinations, as well as all current amplitude and beat frequencies combinations for Ct-Fi and Ct-Lu.

Fig. S6.

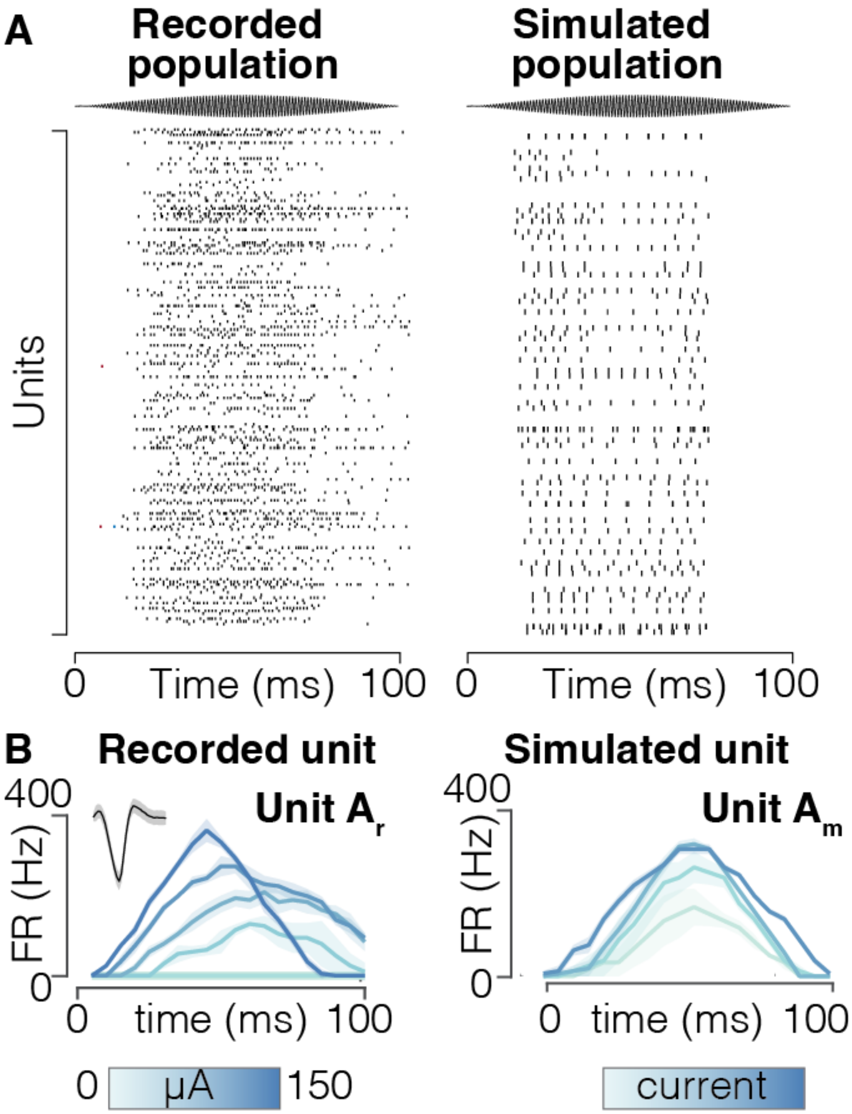


**Figure S6. A biophysically realistic computational model accurately captures the firing dynamics of a neuron population (A)** Raster plot of recorded (left) and simulated (right) population of neurons spiking in response to FAMS stimulation. **(B)** Firing rate profiles over time, averaged across stimulus repetitions, of a recorded (left) and simulated (right) single unit during FAMS stimulation, color coded as a function of stimulus amplitude.
